# Environmental PFOA Exposure Alters Early Developmental Programming during the Maternal–Zygotic Transition

**DOI:** 10.64898/2026.05.21.726952

**Authors:** Zainab Afzal, Vandana Veershetty, Evan Pittman, Charles Hatcher, Deepak Kumar

**Author notes:** Co-first authors (Zainab Afzal, Vandana Veershetty).

## Abstract

Early embryogenesis is governed by precisely timed gene regulatory programs that coordinate cell fate specification, tissue patterning, and morphogenesis. The maternal-to-zygotic transition (MZT) represents a pivotal developmental milestone during which regulatory control shifts from maternally deposited transcripts to activation of the zygotic genome. Disruption of this transition has the potential to alter developmental trajectories with lasting consequences. Per- and polyfluoroalkyl substances (PFAS), environmentally persistent contaminants, have been linked to developmental abnormalities, yet their impact on core embryonic gene regulatory networks especially during MZT is not well understood. Using zebrafish (Danio rerio), a tractable vertebrate model and New Approach Methodology (NAM), we investigated how PFAS exposure during the MZT alters early developmental programming. Embryos were exposed starting at different times within the 8-hour MZT window and collected at 24 hours post-fertilization (hpf) for transcriptomic analysis. Targeted qRT-PCR revealed dysregulation of genes controlling transcriptional activation, lineage specification, proliferation, and differentiation. Whole-transcriptome RNA sequencing (RNA-seq) further identified widespread perturbations in gene networks governing transcriptional regulation, cell signaling, and embryonic morphogenesis. Temporal analysis revealed that exposure beginning at 3.5 hpf, followed by 8 hpf, corresponding to early zygotic genome activation and near completion of zygotic activation, respectively, resulted in the greatest differential gene expression changes. Consistent with these early gene regulatory perturbations, larvae exposed at 8 hpf also exhibited altered behavior at 5 days post-fertilization. Together, these findings demonstrate that PFAS exposure during MZT disrupts the establishment of embryonic gene regulatory networks, linking environmental toxicant exposure to altered developmental patterning and organismal outcomes. This work underscores the vulnerability of early developmental transitions to environmental perturbation and positions MZT as a critical window of susceptibility during development.

## INTRODUCTION

Poly- and perfluoroalkyl substances (PFAS) are a large class of synthetic chemicals widely used in industrial processes and consumer products for more than 60 years due to their surfactant properties and resistance to heat, oil, and water (Buck et al., 2011; kissa, 2001). Although production and use of several long-chain PFAS have declined over the past two decades, their extreme chemical stability has resulted in widespread environmental persistence and ongoing exposure in humans, livestock, and wildlife, with potential consequences for ecosystem and intergenerational health (Bauer et al., 2026; Fenton et al., 2021; Liu et al., 2026; Mellouk et al., 2025; Meng et al., 2022; Satbhai et al., 2025). The strength of the carbon–fluorine bond renders PFAS highly resistant to environmental degradation (Smart BE, 2024), leading to their detection in air, soil, drinking water, and biological samples including human serum and cord blood (Cheng et al., 2025; Falls et al., 2026; Hu et al., 2016; Monroy et al., 2008; Zhai, 2015; Zhi, 2024). Epidemiological and toxicological studies have linked PFAS exposure to hepatotoxicity, endocrine disruption, neurodevelopmental deficits, altered embryonic development, oxidative stress, and carcinogenic outcomes (Gaillard et al., 2025; Kebieche et al., 2025; Luebker et al., 2005; Olsen et al., 2007; Qi et al., 2023; Satbhai et al., 2025). Among PFAS compounds, Perfluorooctanoic acid (PFOA) has been of particular concern due to its historical prevalence in industrial applications and persistence in the environment (Nakayama, 2019; Paul et al., 2009).

To investigate how PFOA exposure affects early vertebrate development and to determine whether specific developmental windows exhibit heightened sensitivity, we utilized embryos of Zebrafish (Danio rerio). Zebrafish embryos are a powerful vertebrate model and a widely used new approach methodology (NAM) (Ball et al., 2025; Kalueff, 2014), for studying the developmental and neurobehavioral impacts of environmental toxicants because of their rapid external development, genetic conservation with humans, optical transparency, and well-characterized behavioral paradigms (Hill et al., 2005; Howe, 2013; Orger & de Polavieja, 2017). As PFAS can bioaccumulate and cross the placental barrier (Gützkow, 2012), exposure during embryonic development particularly concerning. Zebrafish embryos provide a powerful system to study these effects because they develop externally, allowing precise control of exposure concentrations and timing. In addition, large numbers of synchronous embryos can be studied as they undergo a well-defined maternal-to-zygotic transition (MZT) approximately 3–8 hours post-fertilization, during which maternal transcripts are degraded and the zygotic genome becomes transcriptionally active (Giraldez et al., 2006; Schier, 2007; Vastenhouw et al., 2019). This developmental transition provides a tractable system for examining how environmental exposures influence gene regulation during early embryogenesis and for identifying critical windows of exposure vulnerability.

Previous studies have identified transcriptional responses associated with PFAS exposure (Blake et al., 2022; Bline et al., 2025; Britton et al., 2024; Cao et al., 2023; Jantzen et al., 2016; Rericha et al., 2024). To further characterize molecular responses especially during early development, we combined targeted analysis of candidate genes with unbiased transcriptomic profiling to capture both pathway-specific and genome-wide transcriptional changes following PFOA exposure. Using physiologically relevant concentrations of PFOA (Boone, 2019; Stein et al., 2013), we started exposure to zebrafish embryos at multiple timepoints during early developmental windows encompassing the MZT and for all exposed embryos evaluated transcriptional changes at 24 hours post-fertilization, a stage when major organ systems are forming (Kawasaki et al., 2017). Bulk RNA sequencing was used to identify global gene expression changes, disrupted pathways, and regulatory networks associated with developmental toxicity, enabling us to determine which developmental window is most sensitive to PFOA exposure.

To link molecular changes with organismal outcomes, we also integrated transcriptomic analysis with developmental and behavioral assessments. Larval locomotor behavior was quantified at 5 days post-fertilization by measuring distance traveled, swimming velocity, and spatial preference within the recording well. In larval zebrafish, spatial preference within an arena can reflect anxiety-like responses following environmental stimuli. This systems-level approach combining transcriptomic, developmental, and behavioral analyses helps us to move beyond phenotype-based toxicity assessments to identify molecular mechanisms by which PFAS disrupt early vertebrate development. Ultimately, these findings contribute to a deeper understanding of the biological consequences of early-life PFAS exposure and may inform environmental risk assessment and public health strategies.

## MATERIALS AND METHODS

### Animal husbandry of adult Zebrafish

Adult zebrafish (*Danio rerio*, wild-type AB strain) were maintained on a 14:10 h light: dark cycle in embryo media (reverse osmosis water buffered with sea salt, Instant Ocean©) at 27–30 °C in a recirculating system (Aquaneering, San Diego, CA, USA). Water pH was maintained between 7.0 and 7.5. Fish were fed an irradiated diet (Zeigler Bros Inc., Catalog # AH271) twice daily and supplemented with live brine shrimp (EG Artemia, INVE AQUACULTURE Inc.). All zebrafish procedures were approved by the Institutional Animal Care and Use Committee (IACUC) at North Carolina Central University (Protocol number D16-00378; approved 06-01-2022).

### Spawning and embryo collection

Adult zebrafish were bred in a 2:4 male-to-female ratio in spawning tanks separated overnight. At 09:00 the next morning, dividers were removed, and embryos were collected 15 minutes later. Collected embryos were rinsed with reverse osmosis (RO) water and sorted into treatment groups before incubation at 28.5 °C.

### Chemical exposure

Perfluorooctanoic acid (PFOA) (Sigma-Aldrich, Catalog #171468-5G, ≥95% purity) was dissolved in zebrafish embryo medium to prepare a 1000 µg/L stock solution. Working solutions were prepared fresh for each experiment. Embryos from the same clutch were either kept as untreated controls or exposed to 700 ng/L PFOA, a concentration selected to reflect physiologically relevant environmental exposure levels. Control untreated embryos were maintained in embryo medium containing 0.1% methylene blue. For exposure experiments, embryos from the same clutch were sorted into separate Petri dishes and treated with 700 ng/L PFOA in embryo medium 0.1% methylene blue, according to their assigned exposure time. Exposures were initiated at four developmental stages surrounding the maternal-to-zygotic transition (MZT): the 1-cell stage (pre-MZT), 3.5 hours post-fertilization (hpf), 5 hpf, and 8 hpf. Embryos from all treatment groups were collected at 24 hpf for RNA extraction and transcriptomic analysis. For behavioral assays, larvae were maintained in treatment conditions until 5 days post-fertilization (dpf). Chemical exposures were carried out in new Petri dishes (Fisherbrand 100mm Petri Dishes with Clear Lid, Catalog # FB0875713), and treatment solutions were changed daily until behavioral testing at 5 dpf.

### RNA Extraction for pcr and bulk sequencing

At 24 hours post-fertilization (hpf), embryos from each treatment and control group were manually dechorionated, and the same pooling and RNA extraction procedures were followed for both PCR and RNA sequencing. For each sample type, four biological replicates were generated, with each replicate consisting of 20 pooled embryos (n = 4 biological replicates per condition). Each replicate containing 20 embryos was homogenized in 100 µL of TRIzol reagent (Invitrogen, Catalog #15596018). Total RNA was extracted using the Direct-zol™ RNA Miniprep Kit (Zymo Research, Catalog #R2073-A), including an on-column DNase treatment to remove genomic DNA contamination. RNA concentration and purity were assessed using a NanoDrop One spectrophotometer (Thermo Fisher Scientific).

For targeted gene expression analysis, complementary DNA (cDNA) was synthesized from purified RNA. Quantitative real-time PCR (qPCR) was performed to assess the expression of selected genes associated with biological processes including developmental toxicity, stress response, DNA damage, and neuromuscular development. Gene expression levels were normalized to the reference gene *gapdh*, and relative expression changes between PFOA-treated and control groups were calculated using the 2^−ΔΔCt analysis method. Statistical analysis of qRT-PCR data was performed in R. For each target gene, expression levels in the untreated control group were compared independently with each treatment group using two-tailed Student’s *t*-tests. A * was denoted if p < 0.05, ** if p < 0.01, *** if p < 0.001, and “ns” if not statistically significant.

In parallel, for RNA sequencing four biological replicates, purified RNA samples were shipped on dry ice to Novogene for transcriptomic analysis. RNA-seq libraries were prepared using the Illumina TruSeq Stranded mRNA Library Prep protocol and sequenced on the Illumina NovaSeq platform to generate 150-bp paired-end reads. Initial bioinformatics processing included quality control, alignment to the Danio rerio reference genome (GRCz11), and gene-level quantification. Differential gene expression analysis was performed in Novamagic using EdgeR algorithm, with significance thresholds set at an adjusted p-value (false discovery rate, FDR) ≤ 0.05 and an absolute log₂ fold-change ≥ 2.

### Statistical tests and Behavior analysis

Statistical analyses for tail twitch and survival (alive/dead) ratios were performed in R using the ggpubr package. A one-way ANOVA was used to assess overall differences among groups, followed by pairwise Student’s *t*-tests comparing each PFOA exposure group to the untreated control group. P-values for multiple comparisons were adjusted using the Holm method. Statistical significance was defined as p ≤ 0.05.

Neurobehavioral outcomes were assessed at 5 days post-fertilization (dpf) using the DanioVision Observation Chamber (Noldus Information Technology). For assesment, 24 larvae per group, free of visible morphological abnormalities, were placed individually into wells of a 24-well plate. Locomotor activity was recorded using EthoVision XT software across four sequential conditions: no stimulus (60 s), light stimulus (60 s), no stimulus (60 s), and mechanical vibration stimulus (60 s). Total distance moved and zone preference (center vs. periphery) were quantified as measures of locomotor activity and anxiety-like behavior, respectively. All behavioral data was tested using Shapiro-wilk test for normality. When determined to not following a normal distribution, the Kruskal-Wallis test was done to determine if there are significant differences between the five untreated and treatment groups, followed by Dunn post-hoc test to identify specifically which pairwise groups differ significantly. All data plots were visualized in R using the ggplot2 package. A * was denoted if p < 0.05, ** if p < 0.01, *** if p < 0.001, and “ns” if not statistically significant.

## Results

### Embryo Viability and Early Phenotypic Responses to PFOA Exposure

To establish a physiologically relevant exposure dose, we did a literature search through both environmental and human biomonitoring data. A survey of treated water from 25 drinking water treatment plants across the United States reported a median combined PFAS concentration of 19.5 ng/L, with a maximum of 1,100 ng/L (Boone, 2019). PFOA concentrations measured in serum samples from children have reported mean values between 92,500 and 97,300 ng/L (Stein et al., 2013). Based on these benchmarks and a review of the existing literature, an environmentally relevant and within reported contaminated water ranges, 700 ng/L was selected as the exposure dose for all subsequent experiments.

To identify the most vulnerable developmental window, embryos were exposed to PFOA at multiple time points spanning the maternal-to-zygotic transition (MZT), 1-cell (pre-MZT), 3.5 hpf (early blastula), 5 hpf (mid-blastula transition), and 8 hpf (gastrulation onset). In all experimental conditions, PFOA exposure was initiated at different stages within the MZT window, and embryos were allowed to develop and subsequently fixed at 24 hpf (Figure 1A). Viability was assessed at 24 hpf by quantifying the ratio of live to dead embryos, where dead embryos included those exhibiting arrested development or morphological abnormalities. No significant difference in the live-to-dead ratio was observed between PFOA-treated and untreated control groups across any treatment time point (Figure 1B).

**Figure 1.**
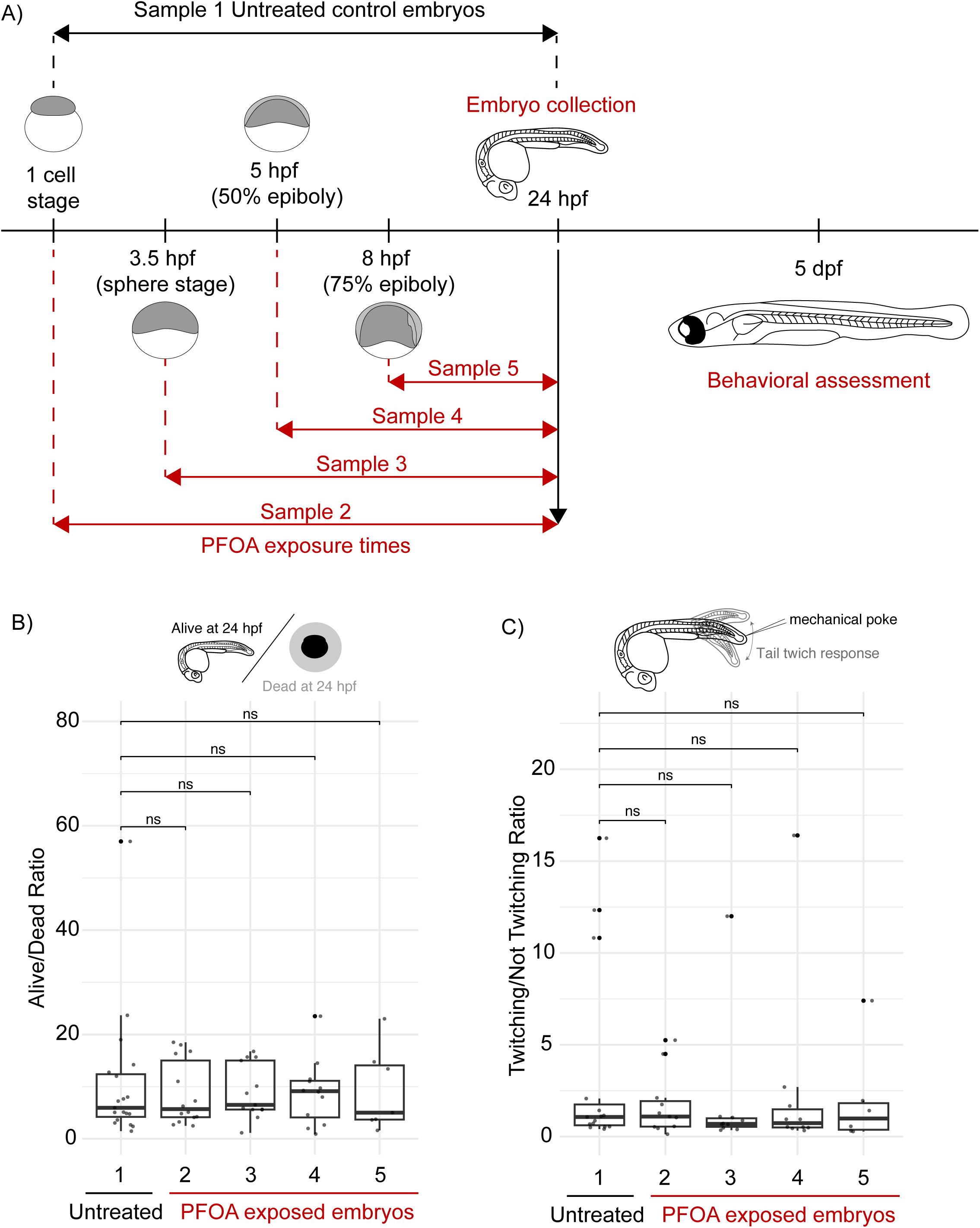
A) Schematic illustrating PFOA exposure time points during the maternal-to-zygotic transition (MZT) and the embryo fixation point at 24 hpf for all subsequent experiments. Embryos were treated at the indicated developmental stages and remained exposed until collection at 24 hpf. B) Box plots to depict embryo viability following PFOA treatment compared to untreated control embryos. C) Box plots to depict manual assessment of tail twitching in response to mechanical stimulation in treated embryos compared to untreated controls (control n = 945; 1-cell treatment n = 578; 3.5 hpf treatment n = 621; 5 hpf treatment n = 228; 8 hpf treatment n = 300). Statistical significance was calculated using using Student’s t-tests with Holm-adjusted p-values for multiple testing correction; p-value <0.05 was considered significant and comparisons were denoted as “ns” where differences were not significant.

At 24 hpf, zebrafish embryos normally exhibit a tail twitch response to mechanical stimulation, reflecting proper neuromuscular junction formation. To assess proper neuromuscular junction formation, each embryo was gently prodded with forceps and scored as responsive or non-responsive by noting whether it twitched or didn’t twitch. Although PFOA-treated embryos showed a trend toward an increased proportion of non-responsive embryos relative to untreated controls, suggesting a potential delay in neuromuscular junction development or formation, this difference did not reach statistical significance (Figure 1C).

### PFOA Exposure Disrupts Signaling Pathways and Transcriptional Programs Essential for Embryonic Development

To assess the molecular consequences of PFOA exposure, we initially analyzed gene expression patterns at 24 hpf using qRT-PCR, focusing on signaling pathways essential for central nervous system, which has been shown to be sensitive to PFOA effect. The Wnt signaling pathway, which plays a critical role in brain development, was significantly disrupted following PFOA exposure, as evidenced by marked decreases in the expression of both *wnt3a* and the Wnt antagonist *sfrp5* (Figure 2A), suggesting a broader perturbation of Wnt pathway regulation rather than a simple loss of pathway activity.

**Figure 2.**
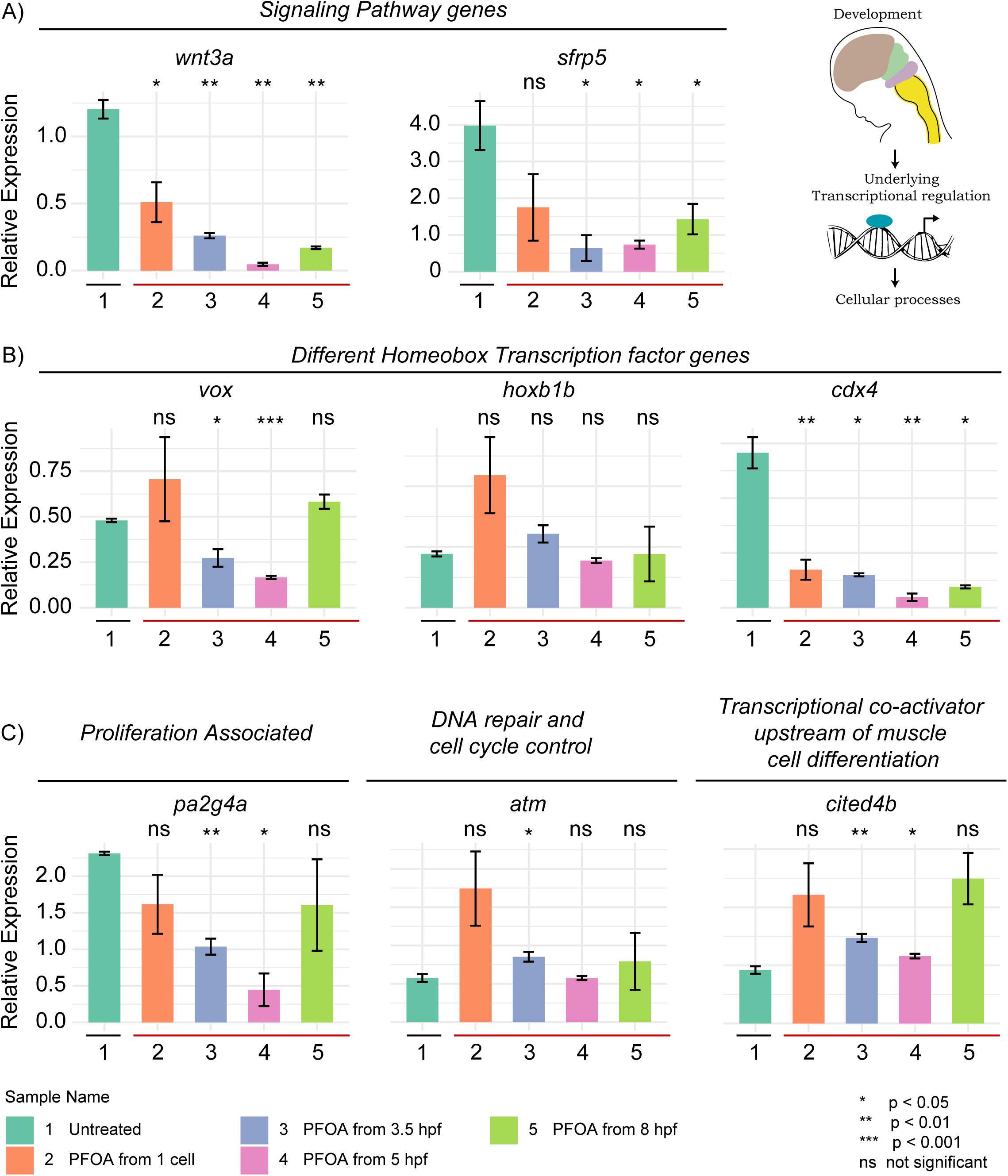
Gene expression changes following PFOA exposure during early development. A) Expression of signaling pathway genes assessed by qRT-PCR at 24 hpf following low-concentration PFOA exposure. Relative gene expression is shown as bar graphs with standard deviation. For each exposure group, N = 20 pooled embryos across five treatment conditions, including a control. B) Expression of selected homeobox transcription factor genes assessed at 24 hpf following PFOA exposure. C) Expression of genes associated with cell proliferation, DNA damage response, and muscle differentiation following PFOA exposure. All statistical analyses were performed using a paired student t-test. Statistical significance is indicated by asterisks: * equals *p* < 0.05, ** equals **p** < 0.01, *** equals ***p*** < 0.001, and “ns” denotes non-significant differences.

Given that Wnt signaling operates upstream of numerous transcriptional programs, that some PFAS compounds can directly bind to DNA (Qin et al., 2024), and that PFAS exposure has been epidemiologically linked to neurodevelopmental deficits (Chen et al., 2026; Wu et al., 2024), we next investigated whether PFOA extends its disruptive effects to transcriptional regulation during early brain development. Towards this, we investigated whether PFOA exposure extends its disruptive effects to transcriptional regulation during early brain development, focusing on homeobox genes which are highly conserved regulators of downstream targets critical for anterior-posterior and dorsal-ventral axis patterning during development (Figure 2B). Expression of *Hoxb1b*, a transcription factor essential for hindbrain segmentation and anterior-posterior patterning of the central nervous system, was not significantly altered by PFOA exposure. In contrast, *Vox*, a homeobox gene involved in dorsal-ventral brain patterning, was significantly downregulated, particularly when exposure began at 3.5 or 5 hpf, indicating a timing-dependent sensitivity to PFOA. *Cdx4*, which functions in posterior body patterning, hindbrain development, and intestinal formation, was consistently downregulated across all PFOA treatment conditions, suggesting that PFOA alters the expression of key homeobox transcription factors in a time-dependent manner, potentially contributing to delayed or abnormal neural development.

To determine whether these transcriptional disruptions also translate into broader cellular dysfunction, we examined the expression of genes involved in cell proliferation, DNA damage response, and cell cycle regulation (Figure 2C). Embryos exposed to PFOA at the 1-cell stage showed no significant changes in these markers; however, exposure initiated at 3.5 hpf resulted in significant reductions in the expression of *pa2g4a*, a proliferation marker, and significant increase in *atm*, a DNA damage response gene, by 24 hpf, suggesting that PFOA exposure during the maternal-to-zygotic transition may impair cellular proliferation and DNA repair mechanisms during a critical window of neural tissue development. Given that these proliferative and DNA repair deficits are likely to have downstream consequences on tissue formation and considering the observed trend toward reduced tail twitch responses in PFOA-treated embryos, a behavior dependent on functional neuromuscular junctions, we further investigated whether these effects extended beyond the nervous system to muscle development. Expression analysis of *cited4b*, a gene involved in muscle cell differentiation, revealed significant upregulation following PFOA exposure, potentially reflecting compensatory or dysregulated neuronal and muscle differentiation signaling programs.

### PFOA Exposure Induces Progressive and Stage-Dependent Transcriptional Disruption Across the Maternal-to-Zygotic Transition

While targeted gene expression analysis provided insight into specific developmental pathways affected by PFOA, we sought to obtain an unbiased, genome-wide view of the transcriptional disruptions occurring across the MZT. To achieve this, bulk RNA sequencing was done on all 5 samples, untreated controls and 4 different treatment times across MZT. The heatmap shows gene expression profiles across samples with red rows being genes overexpressed and blue being genes under expressed genes (Figure 3A). It appears like each stage along MZT is differentially impacted clearly indicative of how PFOA exposure plays into this critical developmental timepoint. Comparing across the PFOA exposure conditions, several gene clusters downregulated in controls are overexpressed, and gene clusters exhibiting higher expression are getting downregulated with exposure. When we compare all PFOA exposure conditions together against untreated controls, Gene Ontology (GO) analysis revealed enrichment across all three ontology categories (Figure 3B). Within biological processes (BP), enriched terms included xenobiotic metabolic processing, cellular responses to abiotic and environmental stimuli, phototransduction, and detection of external stimuli, suggesting that PFOA exposure broadly activates stress response and environmental sensing pathways. Cellular component (CC) terms were enriched for Golgi apparatus-associated structures, including Golgi cisterna and Golgi membrane, as well as condensed chromosomes and the mitochondrial respiratory chain, indicating potential disruptions to intracellular organization and energy metabolism. Molecular function (MF) terms highlighted enrichment in receptor activity, including G-protein coupled photoreceptors and photoreceptor activity, alongside transferase activities such as fucosyltransferase, steroid hydroxylase, and alpha-(1→3)-fucosyltransferase, as well as heme and tetrapyrrole binding and chemokine activity, pointing to disruptions in signal transduction, protein glycosylation, and immune-related molecular functions.

**Figure 3.**
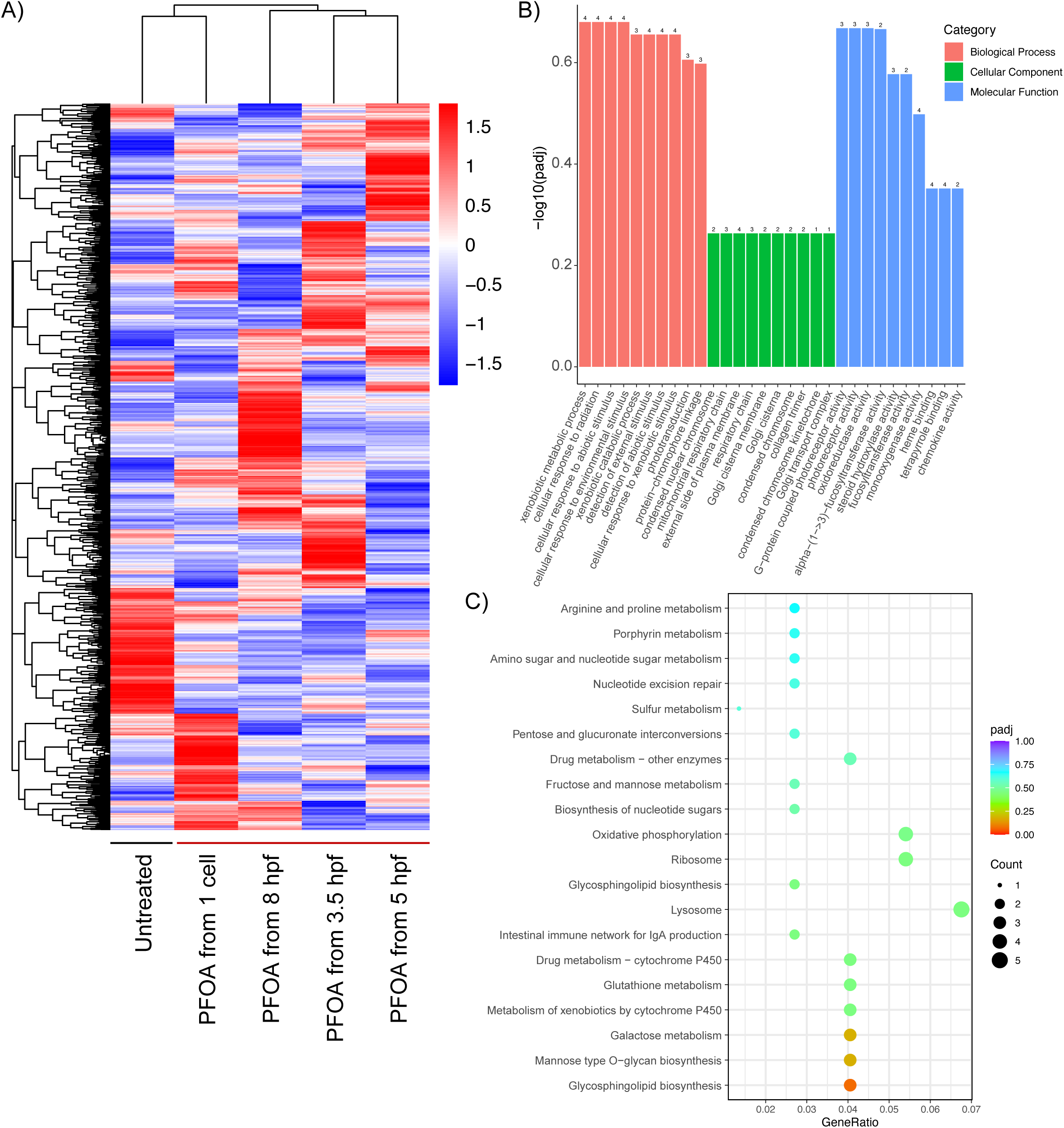
A) Heatmap visualization of global transcriptional changes between untreated and PFOA-treated samples. Gene expression changes were clustered using log2(FPKM+1) values. Red indicates higher expression levels, while blue indicates lower expression levels, with the color gradient reflecting decreasing log2(FPKM+1) values from high to low. B) Bar graph of Gene Ontology (GO) term enrichment analysis of differentially expressed genes (DEGs) identified across all PFOA exposure treatments compared to untreated controls. Enrichment is shown across three categories: Biological Process (BP), Cellular Component (CC), and Molecular Function (MF). C) Dot plot of KEGG pathway enrichment analysis comparing all PFOA exposure treatments to untreated controls, highlighting major pathway disruptions associated with exposure. Dot color represents statistical significance, while dot size indicates the number of DEGs mapped to each KEGG pathway.

Additionally, KEGG pathway analysis revealed enrichment in several functionally related categories (Figure 3C). Metabolic pathways were prominently represented, including amino acid metabolism (arginine and proline metabolism), carbohydrate metabolism (fructose and mannose metabolism, pentose and glucuronate interconversions, amino sugar and nucleotide sugar metabolism), and biosynthesis of nucleotide sugars, suggesting broad disruption of core metabolic processes. Enrichment in oxidative phosphorylation and glutathione metabolism further points to mitochondrial dysfunction and impaired oxidative stress response following PFOA exposure. Notably, multiple xenobiotic and drug metabolism pathways were enriched, including metabolism of xenobiotics by cytochrome P450, drug metabolism via cytochrome P450, and drug metabolism by other enzymes, indicating activation of detoxification machinery in response to PFOA. Glycan biosynthesis pathways were also enriched, including glycosphingolipid biosynthesis and mannose type O-glycan biosynthesis, alongside porphyrin metabolism and nucleotide excision repair, suggesting potential disruptions to lipid signaling, DNA repair, and post-translational protein modification. Finally, enrichment of the intestinal immune network for IgA production pathway suggests that PFOA exposure may also impact immune system development and function as the embryo develops.

Having established the global transcriptional impact of PFOA, we next dissected how these disruptions evolve at each individual time point along MZT to identify stage-specific vulnerabilities. At the 1-cell pre-MZT stage, PFOA induced bidirectional transcriptional changes, with upregulated pathways associated with lipid storage, spliceosomal assembly, and endoderm formation, while pathways governing basal transcription initiation and innate immune complement activation were suppressed, suggesting early metabolic reprogramming and immune dysregulation as an immediate consequence of PFOA exposure. Disruptions to cell cycle progression, DNA damage repair, and oxidative phosphorylation were also evident at this earliest stage, indicating that PFOA begins to interfere with fundamental cellular processes from the very onset of development.

As development progressed to 3.5 and 5 hpf, pathway enrichment patterns suggested the simultaneous activation of compensatory mechanisms alongside ongoing developmental disruption. Wnt signaling showed stage-specific upregulation, potentially reflecting a compensatory response to the Wnt downregulation observed at 24 hpf by qRT-PCR, while DNA damage response and cell proliferation pathways were also upregulated. Conversely, muscle contraction and behavior regulation pathways were suppressed. At 5 hpf specifically, downregulation of miRNA-mediated gene silencing and upregulation of mitotic spindle organization suggest active remodeling of the transcriptional landscape coinciding with the MZT.

By 8 hpf, PFOA exposure produced its most pronounced transcriptional effects, characterized by strong immune activation and broad metabolic suppression. Upregulated pathways included chromatin organization and T cell receptor signaling, reflecting escalating stress-induced epigenetic and immune responses, while ATP metabolism, oxidative phosphorylation, and mitochondrial respiratory chain function were significantly downregulated alongside suppression of Wnt signaling, DNA damage checkpoints, and muscle development programs. These 8 hpf findings are consistent with and directly corroborate the neurodevelopmental and neuromuscular deficits observed at 24 hpf by qRT-PCR, collectively demonstrating that PFOA exposure during the MZT progressively and irreversibly disrupts core transcriptional, metabolic, and developmental programs in a stage-dependent manner.

### Most Critical Time Windows of Exposure During MZT

Comparative analysis of volcano plots across all maternal-to-zygotic transition (MZT) exposure windows revealed that embryos exposed to PFOA beginning at 3.5 hpf exhibited the greatest number of differentially expressed genes relative to untreated controls, with the highest combined total of upregulated and downregulated genes, followed by the 8 hpf exposure group (Figure 4). These findings identify 3.5 hpf and 8 hpf as the most transcriptionally vulnerable stages during the MZT. To further characterize the biological consequences of exposure at each developmental window, Gene Ontology (GO) enrichment analyses were performed across the categories of biological process, cellular component, and molecular function, alongside KEGG pathway analysis to identify disrupted cellular pathways and developmental processes (Supplementary Figures 1–4).

**Figure 4.**
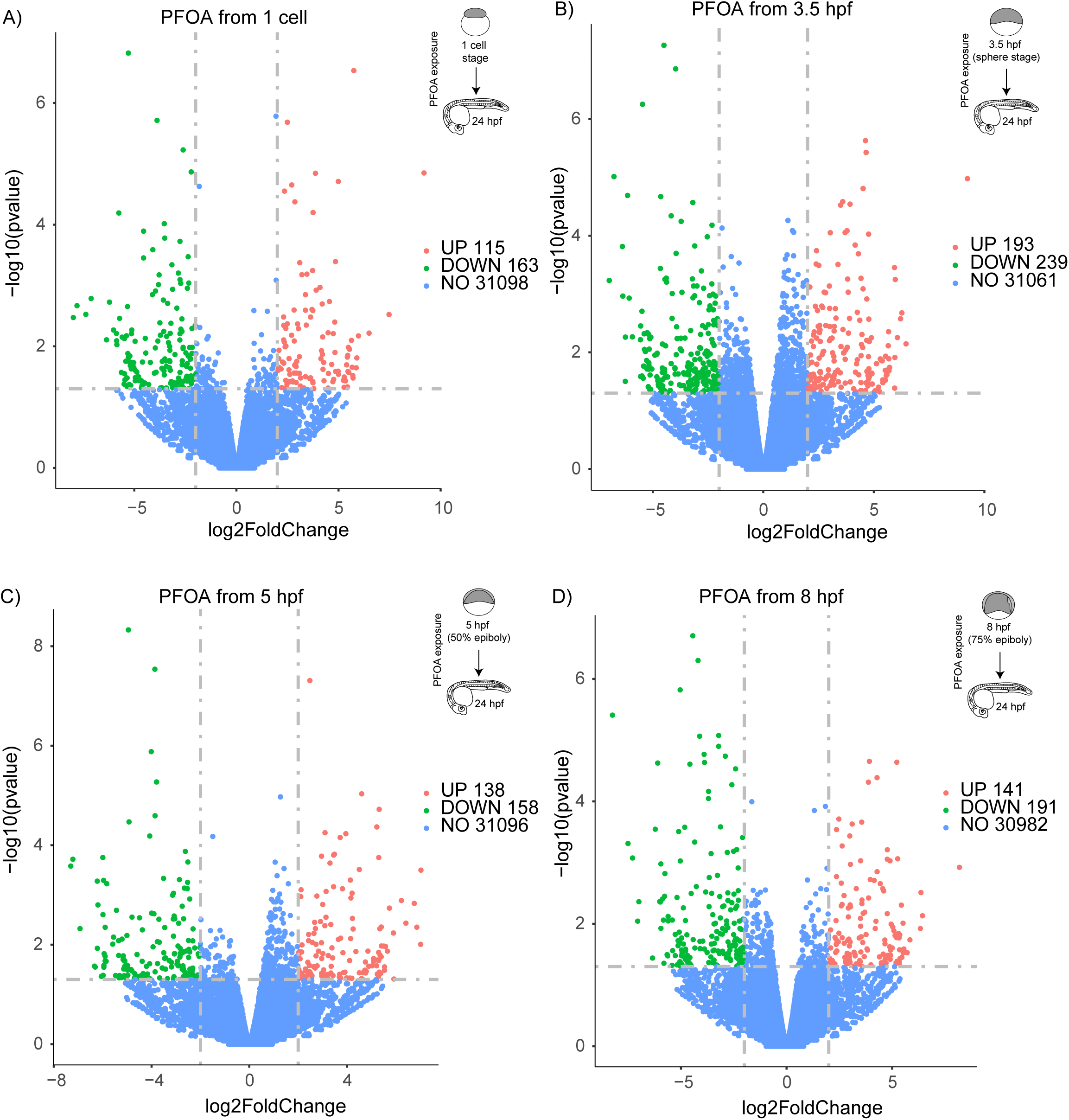
Volcano plots showing differentially expressed genes (DEGs) across all PFOA exposure conditions compared to untreated controls. A) Exposure beginning at the 1-cell stage. B) Exposure beginning at 3.5 hpf. C) Exposure beginning at 5 hpf. D) Exposure beginning at 8 hpf. Non-significant genes are shown in blue, while significantly upregulated genes are shown in red and significantly downregulated genes are shown in green. Significance is defined by a log2 fold change threshold of ≥ 2 (equivalent to a 4-fold change in expression) in either direction relative to untreated controls.

To better understand the molecular basis underlying this heightened sensitivity, GO and KEGG pathway analyses were examined in greater detail for the 3.5 hpf and 8 hpf exposure conditions (Figure 5). Embryos exposed to PFOA beginning at 3.5 hpf showed enrichment of biological processes associated with embryonic development, ion homeostasis, and immune regulation. Cellular component terms indicated disruptions in chromatin organization and RNA processing machinery, including enrichment of nucleosome- and spliceosome-associated components. Molecular function analysis further revealed dysregulation of membrane signaling and ion transport pathways, including chloride channel activity, G protein-coupled receptor signaling, and fucosyltransferase activity (Figure 5A). Consistent with these findings, KEGG pathway analysis identified the spliceosome and p53 signaling pathways as among the most significantly enriched pathways, suggesting that PFOA exposure during this developmental window disrupts RNA processing, genomic stability, and DNA damage response pathways (Supplementary Figure 2B). Additional enrichment of autophagy, oxidative phosphorylation, PPAR signaling, and cardiac muscle contraction pathways further indicates widespread disruption of cellular metabolism and organ-specific developmental programs. Together, these findings suggest that 3.5 hpf represents a particularly sensitive developmental window during which PFOA exposure interferes with core transcriptional, metabolic, and stress-response mechanisms essential for normal embryogenesis.

**Figure 5.**
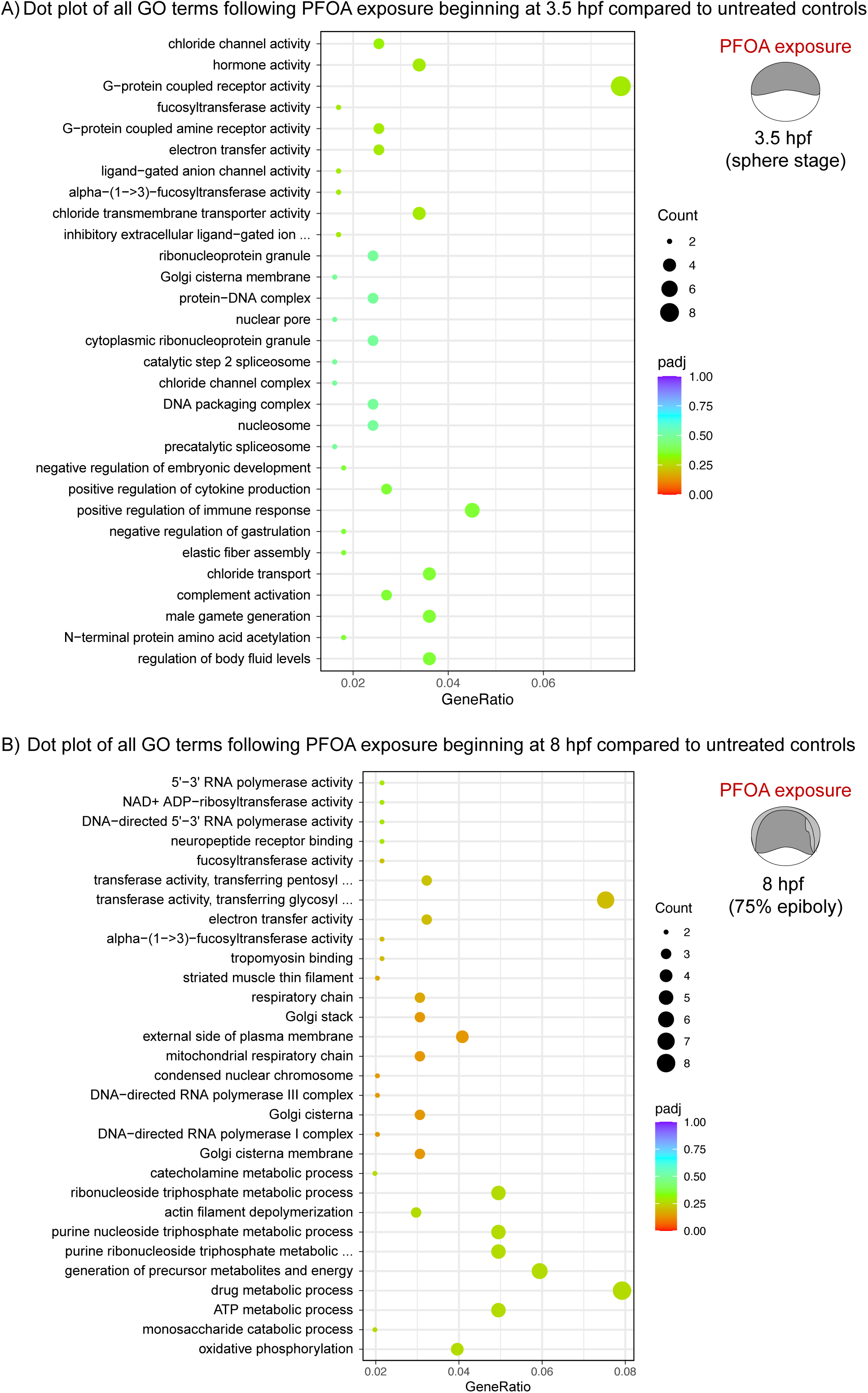
A) Dot plot of of Gene Ontology (GO) term enrichment analysis of differentially expressed genes (DEGs) identified in 3.5 hpf exposure treatment compared to untreated control. Enrichment is shown across three categories: Biological Process, Cellular Component, and Molecular Function. B) Dot plot of of Gene Ontology (GO) term enrichment analysis of differentially expressed genes (DEGs) identified in 8 hpf exposure treatment compared to untreated control. Enrichment is shown across three categories: Biological Process, Cellular Component, and Molecular Function.

In contrast, PFOA exposure beginning at 8 hpf predominantly affected pathways associated with mitochondrial metabolism, energy production, transcriptional regulation, and cellular signaling (Figure 5B; Supplementary Figure 4A). Enriched biological process terms included oxidative phosphorylation, ATP metabolic processes, generation of precursor metabolites and energy, and purine ribonucleoside triphosphate metabolism, indicating substantial disruption of mitochondrial bioenergetics and cellular energy homeostasis. Enrichment of mitochondrial respiratory chain and electron transfer activity terms further supported impairment of oxidative metabolism and mitochondrial function. Additional enrichment of drug metabolic processes and catecholamine metabolism suggested altered stress-response and detoxification pathways following exposure. Cellular component analysis identified significant enrichment of Golgi cisterna membranes, Golgi stacks, condensed nuclear chromosomes, and RNA polymerase I and III complexes, suggesting disruptions in intracellular trafficking, chromatin organization, and transcriptional machinery. Molecular function terms, including tropomyosin binding, actin filament depolymerization, neuropeptide receptor binding, and multiple glycosyltransferase activities, further indicated dysregulation of cytoskeletal organization, signaling pathways, and protein modification processes. KEGG pathway analysis supported these observations, highlighting oxidative phosphorylation and RNA polymerase-associated pathways as major targets of PFOA exposure (Supplementary Figure 4B). Table 1 highlights the top differentially expressed genes identified following PFOA exposure at 8 hpf, many of which are associated with conserved developmental pathways including vesicle trafficking, chromatin regulation, neural development, calcium signaling, and mitochondrial metabolism. Several of these genes are also linked to maternal or early zygotic transcriptional programs during the maternal-to-zygotic transition. Collectively, these findings demonstrate that exposure beginning at 8 hpf disrupts mitochondrial activity, energy metabolism, transcriptional regulation, and developmental signaling pathways during a critical stage of embryonic development.

**Table 1.** Top differentially expressed upregulated and downregulated genes identified by RNA-seq analysis in the 8 hpf PFOA treatment condition, including corresponding human orthologs, conserved developmental functions, and associations with zygotic genome activation during early embryogenesis

| Comparison | Gene | Human Ortholog | Role during Development | MZT Association |
| --- | --- | --- | --- | --- |
| **8 hpf PFOA vs untreated, top upregulated genes** | *ap1m3* | AP1M3 | Protein trafficking between Golgi and endosomes (Chapuy et al., 2008; Shin et al., 2021) | Maternally deposited transcripts (Lee et al., 2013) |
|  | *fam168b* | FAM168B | CNS development, neurite outgrowth, and myelination (Zhang et al., 2020) | Maternal-zygotic transition time ~ 5 hpf (Baia Amaral et al., 2024) |
|  | *cbx8b* | CBX8 | Chromatin organization and progenitor differentiation (Creppe et al., 2014) | Maternally deposited transcripts (Lee et al., 2013) |
|  | *tnfsf14a* | TNFSF14 | Muscle differentiation (Waldemer-Streyer & Chen, 2015) and regulation of neurite growth (Gavalda et al., 2009) | Post-MZT immune activation (Farrell et al., 2018) |
| **8 hpf PFOA vs untreated, top downregulated genes** | *cpne3* | CPNE3 | Calcium-dependent membrane signaling expressed in neutrophil precursors (Cowland et al., 2003) | Maternally deposited transcripts (Lee et al., 2013) |
|  | *p2rx5* | P2RX5 | ATP-gated ion channel with roles in neural, muscle, and cardiac development (Guo et al., 2013; Ruppelt et al., 2001) | Zygotic activation (Lee et al., 2013) |
|  | *mrpl49* | MRPL49 | Mitochondrial ribosomal protein associated with neurodevelopment, hearing function, and energy metabolism (Thomas et al., 2025) | Zygotic activation time ~ 6 hpf (Baia Amaral et al., 2024) |
|  | *uqcrq* | UQCRQ | Embryonic energy metabolism through mitochondrial oxidative phosphorylation (Barel et al., 2008) | Maternal-zygotic transition time ~ 4 hpf (Baia Amaral et al., 2024) |

### PFOA-Induced Disruption Development is Associated with Anxiety-like Behavior at the Larval Stage

The transcriptomic and pathway analyses described above demonstrate that PFOA exposure during the maternal-to-zygotic transition disrupts multiple biological processes critical for normal embryonic development, including mitochondrial metabolism, stress-response signaling, transcriptional regulation, and neurodevelopmental pathways. Given the importance of these processes in nervous system development and organismal homeostasis, we next sought to determine whether these early molecular disruptions resulted in measurable behavioral abnormalities at later developmental stages. To assess the potential long-term functional consequences of embryonic PFOA exposure, larval zebrafish were evaluated for locomotor and anxiety-like behaviors at 5 dpf using the DanioVision tracking system under sequential no-stimulus, light stimulus, and mechanical stimulus conditions.

Heatmap visualizations generated using EthoVision software, in which blue shading depicts regions of movement within the 24-well plate setup, revealed marked increases in both total distance traveled and movement velocity in larvae exposed to PFOA beginning at the 1-cell, 3.5 hpf, 5 hpf, and 8 hpf developmental stages (Figure 6, Supplementary Figures 5–6). Overall, most PFOA-treated larvae exhibited significantly greater locomotor activity compared to untreated controls, consistent with a hyperactivity-like phenotype, with the strongest and most statistically significant effects observed in the 8 hpf exposure group (Figure 7).

**Figure 6.**
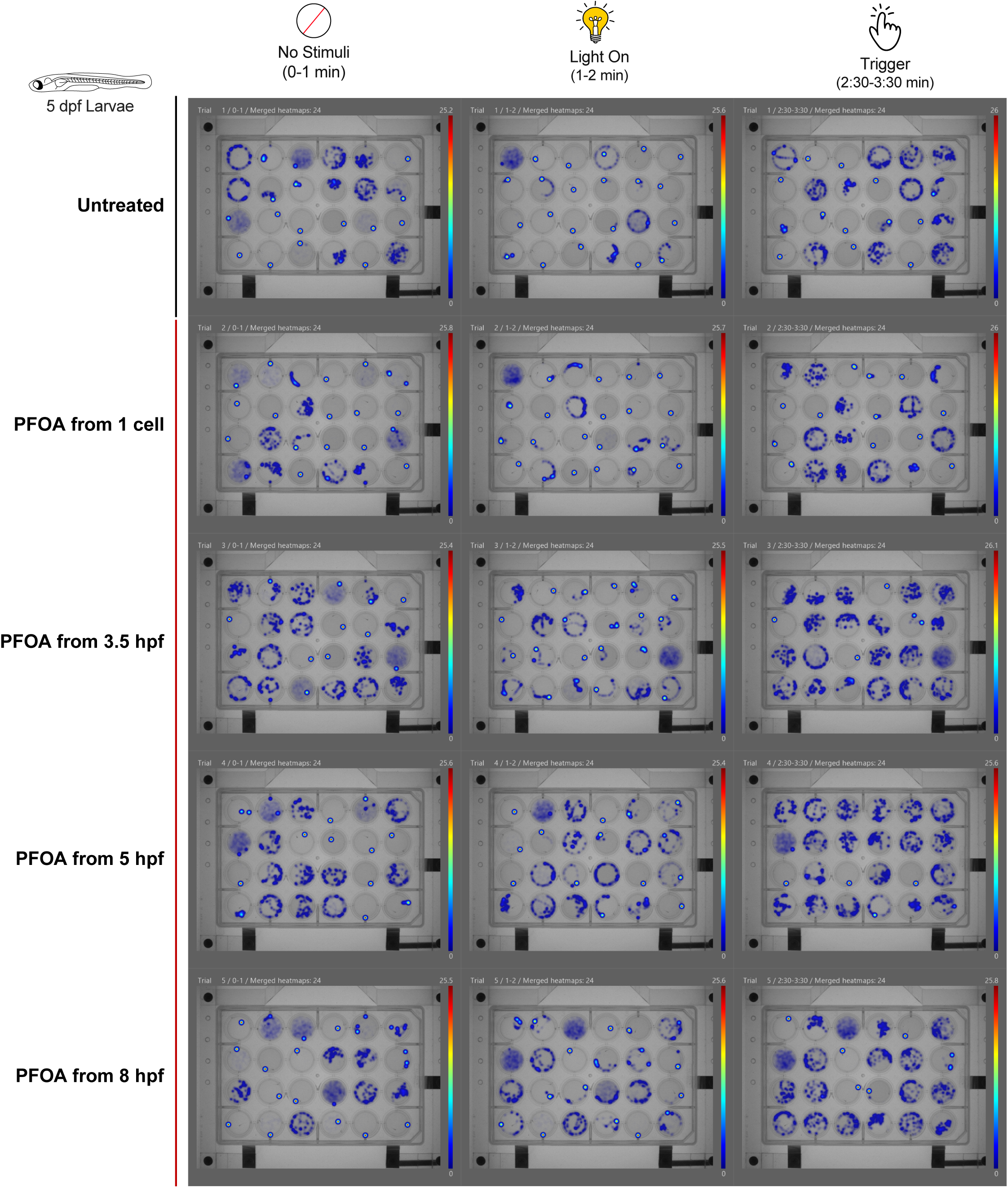
Heatmap visualization of zebrafish larval behavior at 5 dpf under stimulus and no-stimulus conditions. One larva per well was recorded for 3 minutes in a 24-well plate. The heatmap represents the movement patterns and distance traveled by each larva within the well. Blue traces indicate larval movement, while bright puncta with a red center represent locations where the larvae remained stationary for extended periods.

**Figure 7.**
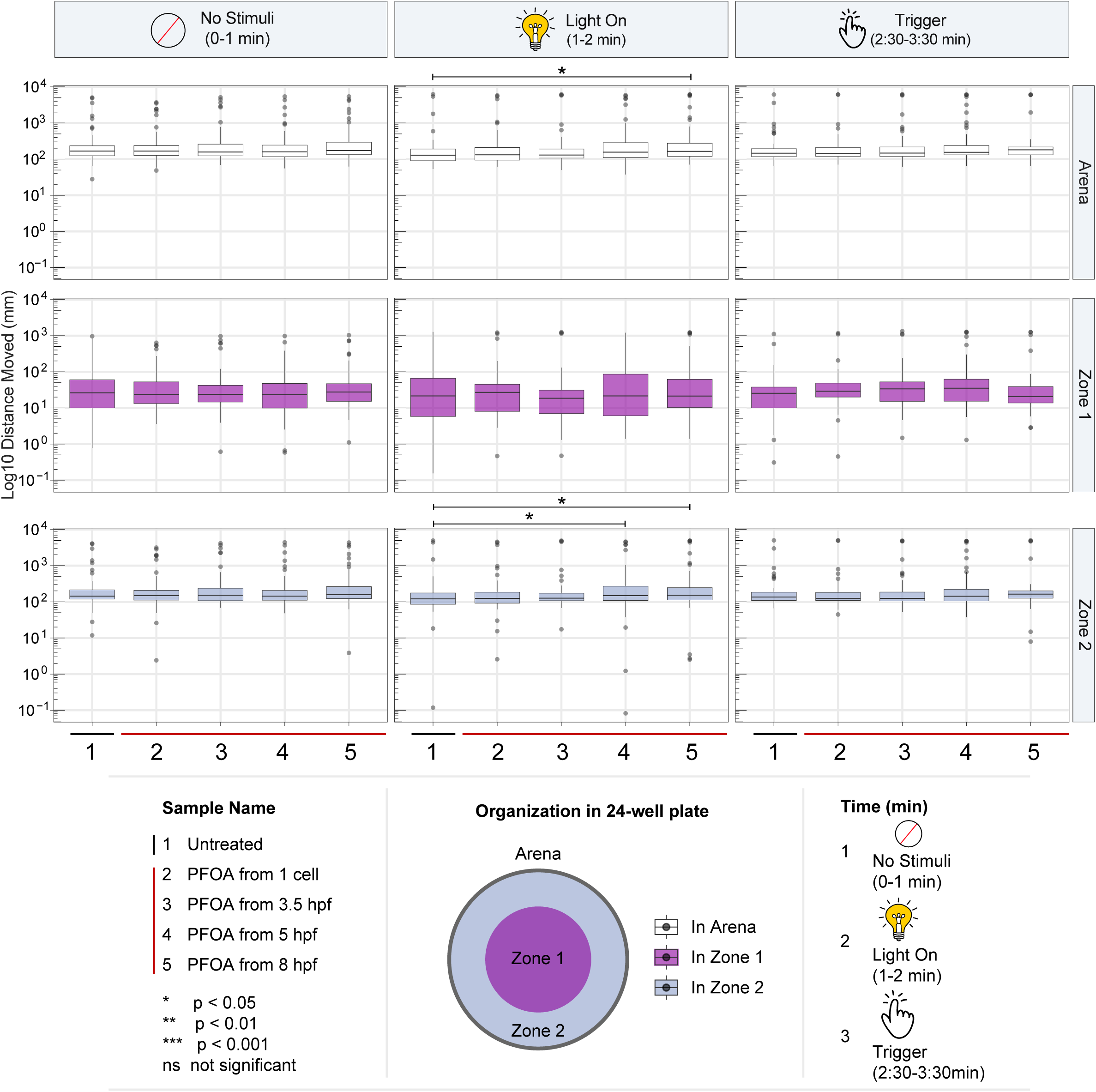
PFOA-treated larvae exhibit anxiety-like behavior in response to stress stimuli. Distance moved by zebrafish larvae (mm) was measured over time (minutes) and analyzed across three regions: Arena (total distance moved), Zone 1 (center), and Zone 2 (periphery). Larvae were individually placed in a 24-well plate (N = 24 per condition). Each well was divided into Zone 1 (center) and Zone 2 (periphery) using EthoVision software, and same boundaries used across all samples. Behavioral activity was recorded across three minutes: 0-1 min - baseline without stimulus, 1-2 min - light stimulus, and 2:30-3:30min - mechanical vibration stimulus. A 30 seconds no stimuli was added between light and trigger stimulation. Box plots show the total distance moved in each area. Asterisks denote statistical significance when comparing distance moved between zone 2 compared to zone 1: * equals *p* < 0.05, ** equals **p** < 0.01, *** equals ***p*** < 0.001, and ns denotes no significant difference in movement.

In addition to increased locomotor activity, PFOA-treated larvae displayed a pronounced preference for the periphery of the well, commonly referred to as “wall-hugging” behavior, relative to untreated larvae. In zebrafish behavioral models, this response is frequently interpreted as an anxiety-like phenotype. Zone analysis further supported this observation, showing significantly greater movement within zone 2, the peripheral region of the well, particularly in larvae exposed beginning at 5 hpf and 8 hpf following light stimulation. The enhanced peripheral swimming behavior observed after light exposure suggests that early embryonic PFOA exposure may alter stress-response and sensory-processing pathways, leading to heightened anxiety-like behavior that persists and can be observed at the larval stage.

## DISCUSSION

This study demonstrates that exposure to perfluorooctanoic acid (PFOA), a persistent and bioaccumulative PFAS compound, disrupts zebrafish development in a stage-dependent manner during the maternal-to-zygotic transition (MZT). Although embryo viability at 24 hpf was not significantly affected, PFOA exposure resulted in multiple developmental and behavioral abnormalities, including trends of reduced tail twitching response, significant transcriptional dysregulation, and anxiety-like locomotor behaviors. These findings suggest that early embryonic exposure to PFOA may perturb key neurodevelopmental processes without causing overt lethality.

Targeted gene expression analysis at 24 hpf showed that PFOA exposure disrupts key developmental signaling pathways. Components of the Wnt pathway, including *wnt3a* and *sfrp5*, were significantly downregulated, suggesting impaired neurodevelopmental signaling. Several homeobox transcription factors involved in embryonic patterning were also affected, with *vox* and *cdx4* reduced across exposure groups, while *hoxb1b* remained unchanged, indicating selective disruption of developmental patterning programs. Genes associated with cell proliferation and stress responses were also altered. *Pa2g4a* expression decreased in embryos exposed at 3.5 and 5 hpf, indicating increased sensitivity of proliferative processes during early development, while the DNA damage response gene *atm* was upregulated. Additionally, increased *cited4b* expression suggests potential compensatory responses or disruption of neuromuscular development following PFOA exposure.

Looking at global changes, transcriptomic profiling further revealed dynamic stage-specific responses to PFOA exposure. The stage-specific transcriptomic differences we observed closely align with the major biological events of the maternal-to-zygotic transition (MZT), suggesting that the developmental timing of PFOA exposure determines which molecular processes are most vulnerable to disruption. At the 1-cell pre-MZT stage, when development relies almost entirely on maternally deposited RNAs and proteins, the observed changes in lipid storage, RNA processing, and oxidative metabolism likely reflect disruption of maternal regulatory and metabolic programs required before zygotic genome activation (ZGA). During the blastula stages (3.5–5 hpf), when embryos undergo ZGA, chromatin remodeling, and maternal transcript clearance, enrichment of DNA damage, RNA processing, spliceosome, and cell proliferation pathways suggests that PFOA interferes with the extensive transcriptional reorganization occurring during MZT, with activation of p53 and stress-response pathways potentially representing compensatory mechanisms to preserve genome integrity. By 8 hpf, after MZT is largely complete and embryos enter early organogenesis, the shift toward metabolic suppression, immune activation, and reduced oxidative phosphorylation suggests functional deficits in mitochondrial activity and cellular homeostasis rather than compensatory transcriptional responses. Because developing tissues at this stage become increasingly dependent on mitochondrial energy production, disruptions in oxidative metabolism may have particularly detrimental effects on tissue differentiation and organ development. Together, these findings indicate that PFOA exposure differentially impacts embryos depending on the developmental processes occurring during each phase of MZT, with early exposure affecting maternal regulatory programs and later exposure impairing zygotic transcription, metabolic maturation, and tissue differentiation.

Behavioral analyses at 5 days post-fertilization further demonstrated that these molecular perturbations translate into functional deficits. PFOA-exposed larvae displayed significantly increased locomotor activity compared to controls, particularly following light stimuli, indicating heightened sensory responsiveness or impaired neural regulation. In addition, treated larvae consistently exhibited thigmotaxis, a wall-hugging behavior commonly interpreted as an anxiety-like phenotype in zebrafish behavioral models. The pronounced peripheral preference following environmental stimuli suggests that early PFOA exposure, especially between 3.5 and 8 hpf, alters neural circuits involved in stress and anxiety responses.

Together, these findings demonstrate that early developmental exposure to PFOA induces coordinated molecular and behavioral changes without causing early lethality. Disruption of key developmental signaling pathways, transcriptional regulators, and metabolic processes likely contributes to the neuromuscular development with behavioral consequences observed at later larval stages. These results provide mechanistic insight into how PFAS exposure during early embryogenesis can alter neurodevelopmental trajectories, consistent with epidemiological studies linking prenatal PFAS exposure to neurobehavioral and metabolic disorders in humans.

## Acknowledgements

This research was supported by the RCMI grant (U5MD012392) awarded to DK, an RCMI Pilot Grant awarded to ZA, and funding from the NC PFAS Collaboratory for the project Molecular Mechanisms of PFAS. We also acknowledge the Julius L. Chambers Biomedical/Biotechnology Research Institute (BBRI) at North Carolina Central University for providing research facilities and resources. We thank the core faculty and staff for their support, and especially Dr. Derek Norford, Dr. Qing Cheng, Dr. Claudia Alberico, and Dr. Alex Gomez. Portions of this work were a part of VV’s Master’s thesis, thus we also acknowledge the Pharmaceutical Sciences Masters’ program, the Pharmaceutical Biomanufacturing Research Institute and Technology Enterprise (BRITE) and thank VV’s committee members, Dr. Kevin Williams and Dr. Vijay Sivaraman, for their guidance and support.

## Figure legends

**Supp Figure 1** A) Bar graph of Gene Ontology (GO) term enrichment analysis of differentially expressed genes (DEGs) identified in 1-cell exposure treatment compared to untreated control. Enrichment is shown across three categories: Biological Process (BP), Cellular Component (CC), and Molecular Function (MF). C) Dot plot of KEGG pathway enrichment analysis identified in 1-cell exposure treatment compared to untreated control. Dot color represents statistical significance, while dot size indicates the number of DEGs mapped to each KEGG pathway.

**Supp Figure 2** A) Bar graph of Gene Ontology (GO) term enrichment analysis of differentially expressed genes (DEGs) identified in 3.5 hpf exposure treatment compared to untreated control. Enrichment is shown across three categories: Biological Process (BP), Cellular Component (CC), and Molecular Function (MF). C) Dot plot of KEGG pathway enrichment analysis identified in 3.5 hpf exposure treatment compared to untreated control. Dot color represents statistical significance, while dot size indicates the number of DEGs mapped to each KEGG pathway.

**Supp Figure 3** A) Bar graph of Gene Ontology (GO) term enrichment analysis of differentially expressed genes (DEGs) identified in 5 hpf exposure treatment compared to untreated control. Enrichment is shown across three categories: Biological Process (BP), Cellular Component (CC), and Molecular Function (MF). C) Dot plot of KEGG pathway enrichment analysis identified in 5 hpf exposure treatment compared to untreated control. Dot color represents statistical significance, while dot size indicates the number of DEGs mapped to each KEGG pathway.

**Supp Figure 4** A) Bar graph of Gene Ontology (GO) term enrichment analysis of differentially expressed genes (DEGs) identified in 8 hpf exposure treatment compared to untreated control. Enrichment is shown across three categories: Biological Process (BP), Cellular Component (CC), and Molecular Function (MF). C) Dot plot of KEGG pathway enrichment analysis identified in 8 hpf exposure treatment compared to untreated control. Dot color represents statistical significance, while dot size indicates the number of DEGs mapped to each KEGG pathway.

**Supp Figure 5** Line graphs depicting the total distance moved by untreated and PFOA-exposed larvae over 3-minute and 30-second time bins. Locomotor activity was measured as total distance traveled in mm within the arena of each well. The arena was divided into two regions: zone 1 (center) and zone 2 (periphery) to assess spatial movement patterns and anxiety-like behavior.

**Supp Figure 6** Line graphs depicting the movement of larvae or their velocity for untreated and PFOA-exposed larvae over 3-minute and 30-second time bins. Locomotor activity was measured as total distance traveled in mm over time in seconds (mm/s) within the arena of each well. The arena was divided into two regions: zone 1 (center) and zone 2 (periphery) to assess spatial movement patterns and anxiety-like behavior.

## References

Baia Amaral, D., Egidy, R., Perera, A., & Bazzini, A. A. (2024). miR-430 regulates zygotic mRNA during zebrafish embryogenesis. Genome Biol, 25(1), 74. 10.1186/s13059-024-03197-8

Ball, J. S., Tochwin, A., Winter, M. J., Trznadel, M., Currie, R., Wolton, K., French, J. M., Hetheridge, M. J., & Tyler, C. R. (2025). Determination of the zebrafish embryo developmental toxicity assessment (ZEDTA) as an alternative non-mammalian approach for the safety assessment of agrochemicals. Reprod Toxicol, 132, 108837. 10.1016/j.reprotox.2025.108837

Barel, O., Shorer, Z., Flusser, H., Ofir, R., Narkis, G., Finer, G., Shalev, H., Nasasra, A., Saada, A., & Birk, O. S. (2008). Mitochondrial complex III deficiency associated with a homozygous mutation in UQCRQ. Am J Hum Genet, 82(5), 1211–1216. 10.1016/j.ajhg.2008.03.020

Bauer, R. A., Yang, Z., Petriello, M., Zhang, S., Stapleton, H. M., Adgate, J. L., & Carignan, C. C. (2026). Associations of serum PFAS with COVID-19 antibody levels among fully vaccinated adults. Environ Res, 298, 124154. 10.1016/j.envres.2026.124154

Blake, B. E., Miller, C. N., Nguyen, H., Chappell, V. A., Phan, T. P., Phadke, D. P., Balik-Meisner, M. R., Mav, D., Shah, R. R., & Fenton, S. E. (2022). Transcriptional pathways linked to fetal and maternal hepatic dysfunction caused by gestational exposure to perfluorooctanoic acid (PFOA) or hexafluoropropylene oxide-dimer acid (HFPO-DA or GenX) in CD-1 mice. Ecotoxicol Environ Saf, 248, 114314. 10.1016/j.ecoenv.2022.114314

Bline, A. P., Jiang, H., Levenson, M., & Allard, P. (2025). A systems toxicology approach implicates post-transcriptional regulatory networks in reproductive defects from PFAS exposure. Toxicol Sci, 208(1), 61–81. 10.1093/toxsci/kfaf111

Boone, J. S. (2019). Per- and polyfluoroalkyl substances in source and treated drinking waters of the United States. 10.1016/j.scitotenv.2018.10.245

Britton, K. N., Judson, R. S., Hill, B. N., Jarema, K. A., Olin, J. K., Knapp, B. R., Lowery, M., Feshuk, M., Brown, J., & Padilla, S. (2024). Using Zebrafish to Screen Developmental Toxicity of Per- and Polyfluoroalkyl Substances (PFAS). Toxics, 12(7). 10.3390/toxics12070501

Buck, R. C., Franklin, J., Berger, U., Conder, J. M., Cousins, I. T., de Voogt, P., Jensen, A. A., Kannan, K., Mabury, S. A., & van Leeuwen, S. P. (2011). Perfluoroalkyl and polyfluoroalkyl substances in the environment: terminology, classification, and origins. Integr Environ Assess Manag, 7(4), 513–541. 10.1002/ieam.258

Cao, W., Horzmann, K., Schemera, B., Petrofski, M., Kendall, T., Spooner, J., Rynders, P. E., VandeBerg, J. L., & Wang, X. (2023). Blood transcriptome responses to PFOA and GenX treatment in the marsupial biomedical model Monodelphis domestica. Front Genet, 14, 1073461. 10.3389/fgene.2023.1073461

Chapuy, B., Tikkanen, R., Muhlhausen, C., Wenzel, D., von Figura, K., & Honing, S. (2008). AP-1 and AP-3 mediate sorting of melanosomal and lysosomal membrane proteins into distinct post-Golgi trafficking pathways. Traffic, 9(7), 1157–1172. 10.1111/j.1600-0854.2008.00745.x

Chen, W. R., Huang, Y., Zhou, W., Yang, J., Ren, T., Zhang, L., Zhu, T., Shan, X., Du, Y., Zhou, G., Liu, Y., Sun, Y., Zhang, Q., Li, W. G., Dong, Y., & Li, F. (2026). Human cerebral organoids reveal PFOA-induced axonal injury as a conserved mechanism of neurodevelopmental disruption. Environ Int, 210, 110233. 10.1016/j.envint.2026.110233

Cheng, L., Teagle, S., Enders, J. R., Weed, R. A., Nichols, H. B., Knappe, D. R. U., & Hoppin, J. A. (2025). Historical Blood Serum Samples from Wilmington, North Carolina: The Importance of Ultrashort-Chain Per- and Polyfluoroalkyl Substances. Environ Sci Technol, 59(43), 23125–23135. 10.1021/acs.est.5c08146

Cowland, J. B., Carter, D., Bjerregaard, M. D., Johnsen, A. H., Borregaard, N., & Lollike, K. (2003). Tissue expression of copines and isolation of copines I and III from the cytosol of human neutrophils. J Leukoc Biol, 74(3), 379–388. 10.1189/jlb.0203083

Creppe, C., Palau, A., Malinverni, R., Valero, V., & Buschbeck, M. (2014). A Cbx8-containing polycomb complex facilitates the transition to gene activation during ES cell differentiation. PLoS Genet, 10(12), e1004851. 10.1371/journal.pgen.1004851

Falls, A. T., Boatman, A. K., Ryan, J. P., Solosky, A. M., Dodds, J. N., Chappel, J. R., Fry, A. N., Kirkwood-Donelson, K. I., Stapleton, H. M., & Baker, E. S. (2026). Increasing PFAS concentrations in human serum correlate with elevated blood lipid levels. Env Sci Adv, 5(3), 885–899. 10.1039/d5va00483g

Fenton, S. E., Ducatman, A., Boobis, A., DeWitt, J. C., Lau, C., Ng, C., Smith, J. S., & Roberts, S. M. (2021). Per- and Polyfluoroalkyl Substance Toxicity and Human Health Review: Current State of Knowledge and Strategies for Informing Future Research. Environ Toxicol Chem, 40(3), 606–630. 10.1002/etc.4890

Gaillard, L., Barouki, R., Blanc, E., Coumoul, X., & Andreau, K. (2025). Per- and polyfluoroalkyl substances as persistent pollutants with metabolic and endocrine-disrupting impacts. Trends Endocrinol Metab, 36(3), 249–261. 10.1016/j.tem.2024.07.021

Gavalda, N., Gutierrez, H., & Davies, A. M. (2009). Developmental regulation of sensory neurite growth by the tumor necrosis factor superfamily member LIGHT. J Neurosci, 29(6), 1599–1607. 10.1523/JNEUROSCI.3566-08.2009

Giraldez, A. J., Mishima, Y., Rihel, J., Grocock, R. J., Van Dongen, S., Inoue, K., Enright, A. J., & Schier, A. F. (2006). Zebrafish MiR-430 promotes deadenylation and clearance of maternal mRNAs. Science, 312(5770), 75–79. 10.1126/science.1122689

Guo, W., Zhang, Z., Liu, X., Burnstock, G., Xiang, Z., & He, C. (2013). Developmental expression of P2X5 receptors in the mouse prenatal central and peripheral nervous systems. Purinergic Signal, 9(2), 239–248. 10.1007/s11302-012-9346-z

Gützkow, K. B. (2012). Placental transfer of perfluorinated compounds is selective – A Norwegian Mother and Child sub-cohort study. 10.1016/j.ijheh.2011.08.011

Hill, A. J., Teraoka, H., Heideman, W., & Peterson, R. E. (2005). Zebrafish as a model vertebrate for investigating chemical toxicity. Toxicol Sci, 86(1), 6–19. 10.1093/toxsci/kfi110

Howe, K. (2013). The zebrafish reference genome sequence and its relationship to the human genome. Nature, 496, 498–503. 10.1038/nature12111

Hu, X. C., Andrews, D. Q., Lindstrom, A. B., Bruton, T. A., Schaider, L. A., Grandjean, P., Lohmann, R., Carignan, C. C., Blum, A., Balan, S. A., Higgins, C. P., & Sunderland, E. M. (2016). Detection of Poly- and Perfluoroalkyl Substances (PFASs) in U.S. Drinking Water Linked to Industrial Sites, Military Fire Training Areas, and Wastewater Treatment Plants. Environ Sci Technol Lett, 3(10), 344–350. 10.1021/acs.estlett.6b00260

Jantzen, C. E., Annunziato, K. M., & Cooper, K. R. (2016). Behavioral, morphometric, and gene expression effects in adult zebrafish (Danio rerio) embryonically exposed to PFOA, PFOS, and PFNA. Aquat Toxicol, 180, 123–130. 10.1016/j.aquatox.2016.09.011

Kalueff, A. V. (2014). Zebrafish as an emerging model for studying complex brain disorders. Trends in Pharmacological Sciences, 35(2), 63–75. 10.1016/j.tips.2013.12.002

Kawasaki, T., Maeno, A., Shiroishi, T., & Sakai, N. (2017). Development and growth of organs in living whole embryo and larval grafts in zebrafish. Sci Rep, 7(1), 16508. 10.1038/s41598-017-16642-5

Kebieche, N., Yim, S., Lambert, C., & Soulimani, R. (2025). Epigenetic and Genotoxic Mechanisms of PFAS-Induced Neurotoxicity: A Molecular and Transgenerational Perspective. Toxics, 13(8). 10.3390/toxics13080629

kissa, E. (2001). Kissa E.Fluorinated surfactants and repellents, 2nd ed.; CRC Press: Boca Raton, FL, 2001. Kissa E.Fluorinated surfactants and repellents, 2nd ed.; CRC Press: Boca Raton, FL, 2001.

Lee, M. T., Bonneau, A. R., Takacs, C. M., Bazzini, A. A., DiVito, K. R., Fleming, E. S., & Giraldez, A. J. (2013). Nanog, Pou5f1 and SoxB1 activate zygotic gene expression during the maternal-to-zygotic transition. Nature, 503(7476), 360-364. 10.1038/nature12632

Liu, H., Kress, A. M., Yu, E. X., Ning, X., Ghassabian, A., Kahn, L. G., Mehta-Lee, S., Brubaker, S., Alshawabkeh, A., Meeker, J., Camargo, C. A., Jr., Suglia, S. F., Elliott, A. J., Ferrara, A., Zhu, Y., Gern, J. E., Bendixsen, C., Gold, D. R., Cassidy-Bushrow, A. E.,…Cohorts. (2026). Racial and ethnic disparities in environmental chemical exposures and hypertensive disorders of pregnancy: The ECHO-wide cohort study. Environ Pollut, 392, 127452. 10.1016/j.envpol.2025.127452

Luebker, D. J., York, R. G., Hansen, K. J., Moore, J. A., & Butenhoff, J. L. (2005). Neonatal mortality from in utero exposure to perfluorooctanesulfonate (PFOS) in Sprague-Dawley rats: dose-response, and biochemical and pharamacokinetic parameters. Toxicology, 215(1-2), 149–169. 10.1016/j.tox.2005.07.019

Mellouk, N., Marchese, M. J., Gao, F., Liang, S., & Feng, L. (2025). Effects of Perfluorobutane Sulfonate (PFBS) on Female Reproduction, Pregnancy, and Birth Outcomes. Obstet Gynecol Surv, 80(10), 657–672. 10.1097/OGX.0000000000001440

Meng, P., DeStefano, N. J., & Knappe, D. R. U. (2022). Extraction and Matrix Cleanup Method for Analyzing Novel Per- and Polyfluoroalkyl Ether Acids and Other Per- and Polyfluoroalkyl Substances in Fruits and Vegetables. Journal of Agricultural and Food Chemistry, 70(16), 4792–4804. 10.1021/acs.jafc.1c07665

Monroy, R., Morrison, K., Teo, K., Atkinson, S., Kubwabo, C., Stewart, B., & Foster, W. G. (2008). Serum levels of perfluoroalkyl compounds in human maternal and umbilical cord blood samples. Environ Res, 108(1), 56–62. 10.1016/j.envres.2008.06.001

Nakayama, S. F. (2019). Worldwide trends in tracing poly- and perfluoroalkyl substances (PFAS) in the environment. 10.1016/j.trac.2019.02.011

Olsen, G. W., Burris, J. M., Ehresman, D. J., Froehlich, J. W., Seacat, A. M., Butenhoff, J. L., & Zobel, L. R. (2007). Half-life of serum elimination of perfluorooctanesulfonate,perfluorohexanesulfonate, and perfluorooctanoate in retired fluorochemical production workers. Environ Health Perspect, 115(9), 1298–1305. 10.1289/ehp.10009

Orger, M. B., & de Polavieja, G. G. (2017). Zebrafish Behavior: Opportunities and Challenges. Annu Rev Neurosci, 40, 125–147. 10.1146/annurev-neuro-071714-033857

Paul, A. G., Jones, K. C., & Sweetman, A. J. (2009). A first global production, emission, and environmental inventory for perfluorooctane sulfonate. Environ Sci Technol, 43(2), 386–392. 10.1021/es802216n

Qi, Q., Niture, S., Gadi, S., Arthur, E., Moore, J., Levine, K. E., & Kumar, D. (2023). Per- and polyfluoroalkyl substances activate UPR pathway, induce steatosis and fibrosis in liver cells. Environ Toxicol, 38(1), 225–242. 10.1002/tox.23680

Qin, C., Xiang, L., Wang, Y. Z., Yu, P. F., Meng, C., Li, Y. W., Zhao, H. M., Hu, X., Gao, Y., & Mo, C. H. (2024). Binding interaction of environmental DNA with typical emerging perfluoroalkyl acids and its impact on bioavailability. Sci Total Environ, 906, 167392. 10.1016/j.scitotenv.2023.167392

Rericha, Y., St Mary, L., Truong, L., McClure, R., Martin, J. K., Leonard, S. W., Thunga, P., Simonich, M. T., Waters, K. M., Field, J. A., & Tanguay, R. L. (2024). Diverse PFAS produce unique transcriptomic changes linked to developmental toxicity in zebrafish. Front Toxicol, 6, 1425537. 10.3389/ftox.2024.1425537

Ruppelt, A., Ma, W., Borchardt, K., Silberberg, S. D., & Soto, F. (2001). Genomic structure, developmental distribution and functional properties of the chicken P2X(5) receptor. J Neurochem, 77(5), 1256–1265. 10.1046/j.1471-4159.2001.00348.x

Satbhai, K. M., Marques, E. S., Ranjan, R., & Timme-Laragy, A. R. (2025). Single-cell RNA sequencing reveals tissue-specific transcriptomic changes induced by perfluorooctanesulfonic acid (PFOS) in larval zebrafish (Danio rerio). J Hazard Mater, 489, 137515. 10.1016/j.jhazmat.2025.137515

Schier, A. F. (2007). The maternal-zygotic transition: death and birth of RNAs. Science, 316(5823), 406–407. 10.1126/science.1140693

Shin, J., Nile, A., & Oh, J. W. (2021). Role of adaptin protein complexes in intracellular trafficking and their impact on diseases. Bioengineered, 12(1), 8259–8278. 10.1080/21655979.2021.1982846

Smart BE, D. D. (2024). Proposal for Chemical Class Prohibition:

Per- and Polyfluoroalkyl Substances (PFAS)

Prohibited in Paints & Coatings, Cleaning & Degreasing

Agents, Adhesives, and Floor-Care Products. https://greenseal.org/wp-content/uploads/PFAS_Prohibition_Standard_Revision_Proposal-November_2024.pdf

Stein, C. R., Savitz, D. A., & Bellinger, D. C. (2013). Perfluorooctanoate and neuropsychological outcomes in children. Epidemiology, 24(4), 590–599. 10.1097/EDE.0b013e3182944432

Thomas, H. B., Demain, L. A. M., Cabrera-Orefice, A., Schrauwen, I., Shamseldin, H. E., Rea, A., Bharadwaj, T., Smith, T. B., Olahova, M., Thompson, K., He, L., Kaur, N., Shukla, A., Abukhalid, M., Ansar, M., Rehman, S., Riazuddin, S., Abdulwahab, F., Smith, J. M.,…Newman, W. G. (2025). Bi-allelic variants in MRPL49 cause variable clinical presentations, including sensorineural hearing loss, leukodystrophy, and ovarian insufficiency. Am J Hum Genet, 112(4), 952–962. 10.1016/j.ajhg.2025.02.005

Vastenhouw, N. L., Cao, W. X., & Lipshitz, H. D. (2019). The maternal-to-zygotic transition revisited. Development, 146(11). 10.1242/dev.161471

Waldemer-Streyer, R. J., & Chen, J. (2015). Myocyte-derived Tnfsf14 is a survival factor necessary for myoblast differentiation and skeletal muscle regeneration. Cell Death Dis, 6(12), e2026. 10.1038/cddis.2015.375

Wu, S., Xie, J., Zhao, H., Zhao, X., Sanchez, O. F., Rochet, J. C., Freeman, J. L., & Yuan, C. (2024). Developmental neurotoxicity of PFOA exposure on hiPSC-derived cortical neurons. Environ Int, 190, 108914. 10.1016/j.envint.2024.108914

Zhai, Z. (2015). A 17-fold increase of trifluoroacetic acid in landscape waters of Beijing, China during the last decade. Chemosphere, 129, 110–117. 10.1016/j.chemosphere.2014.09.033

Zhang, T., Guan, P., Liu, W., Zhao, G., Fang, Y., Fu, H., Gui, J. F., Li, G., & Liu, J. X. (2020). Copper stress induces zebrafish central neural system myelin defects via WNT/NOTCH-hoxb5b signaling and pou3f1/fam168a/fam168b DNA methylation. Biochim Biophys Acta Gene Regul Mech, 1863(10), 194612. 10.1016/j.bbagrm.2020.194612

Zhi, Y. (2024). Environmental occurrence and biotic concentrations of ultrashort-chain perfluoroalkyl acids: Overlooked global organofluorine contaminants. Environmental Science & Technology, 58(49), 21393–21410. 10.1021/acs.est.4c04453

