## Supplementary material for "Environmental PFOA Exposure Alters Early Developmental Programming during the Maternal–Zygotic Transition": Supp Figure

**Supp Figure 1**

A) GO term analysis for PFOA exposure from 1 cell compared against untreated

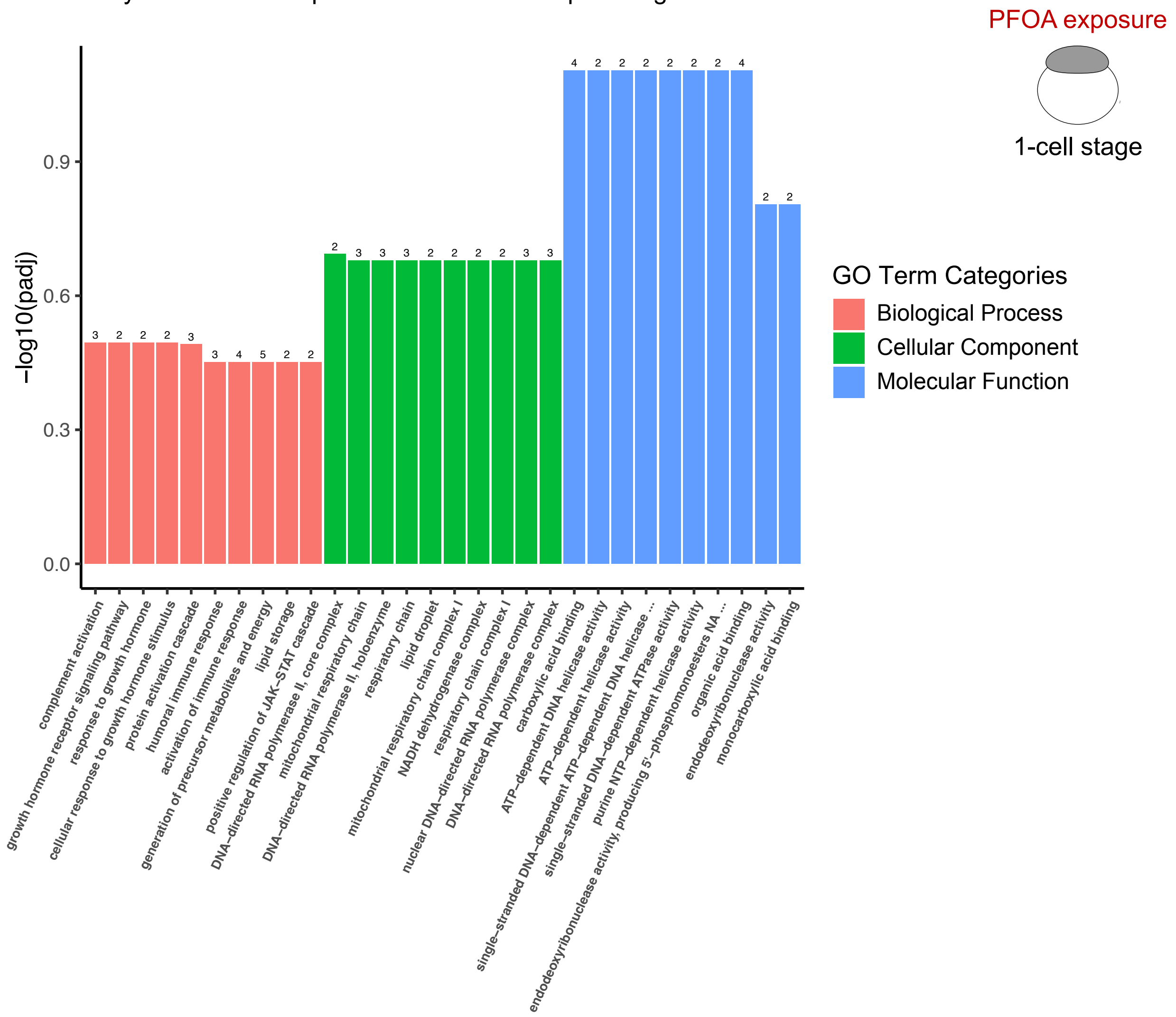


B) Kegg pathway analysis for PFOA exposure from 1 cell compared against untreated

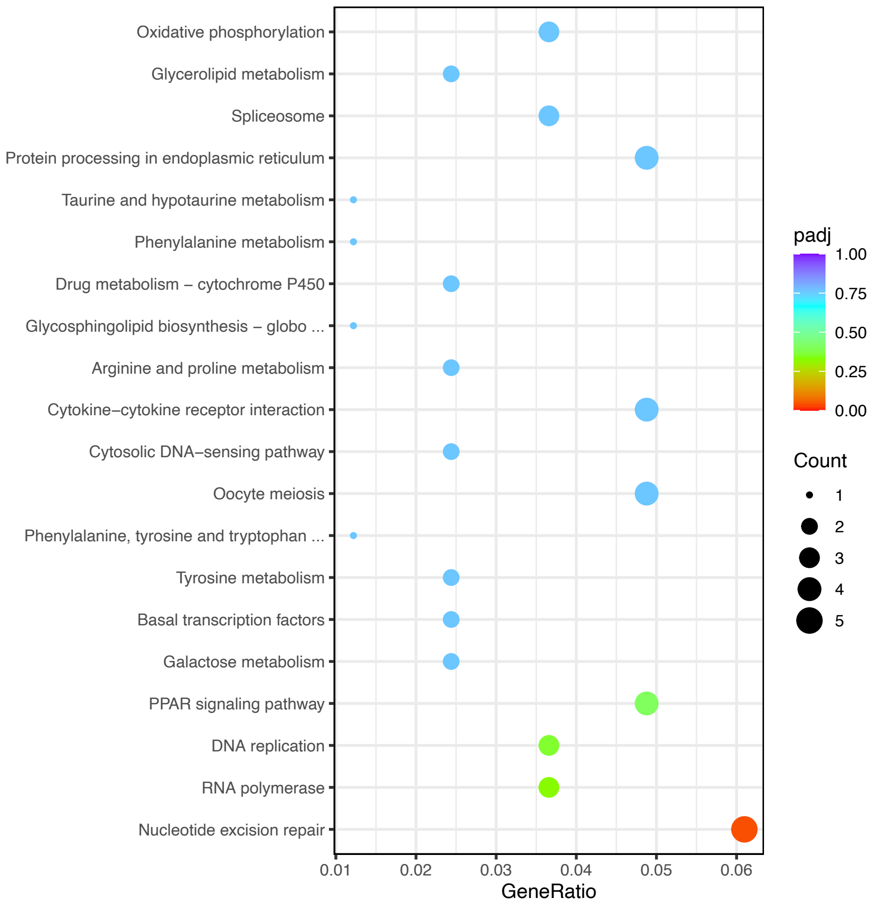

**Supp Figure 2**

A) GO term analysis for PFOA exposure from 3.5hpf compared against untreated separated by categories

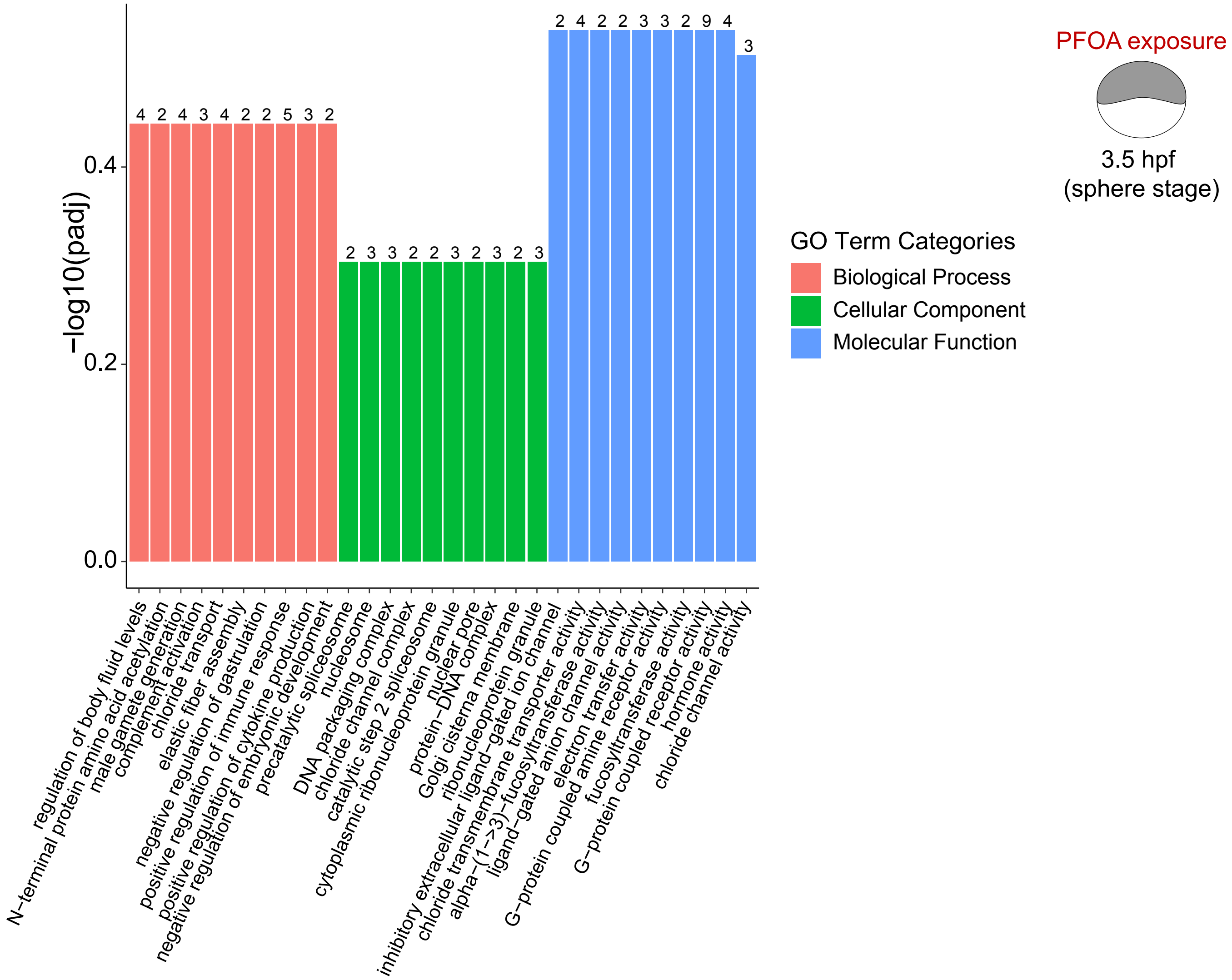


B) Kegg pathway analysis for PFOA exposure from 3.5hpf compared against untreated

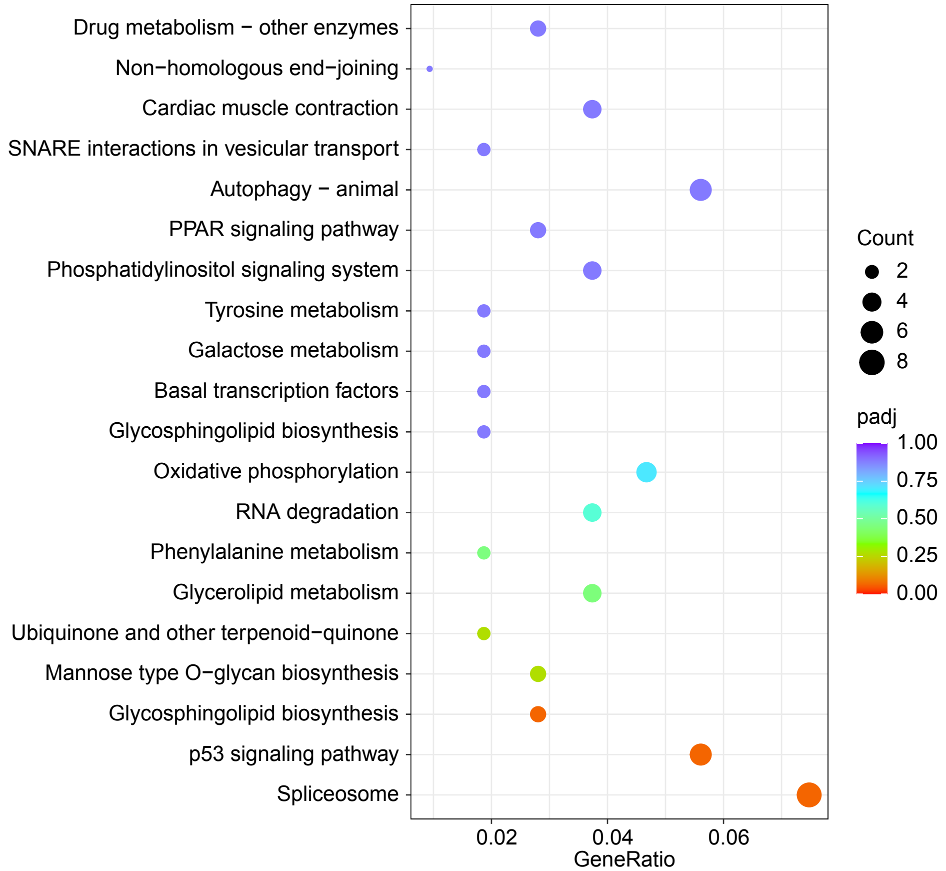

**Supp Figure 3**

A) GO term analysis for PFOA exposure from 5 hpf compared against untreated separated by categories

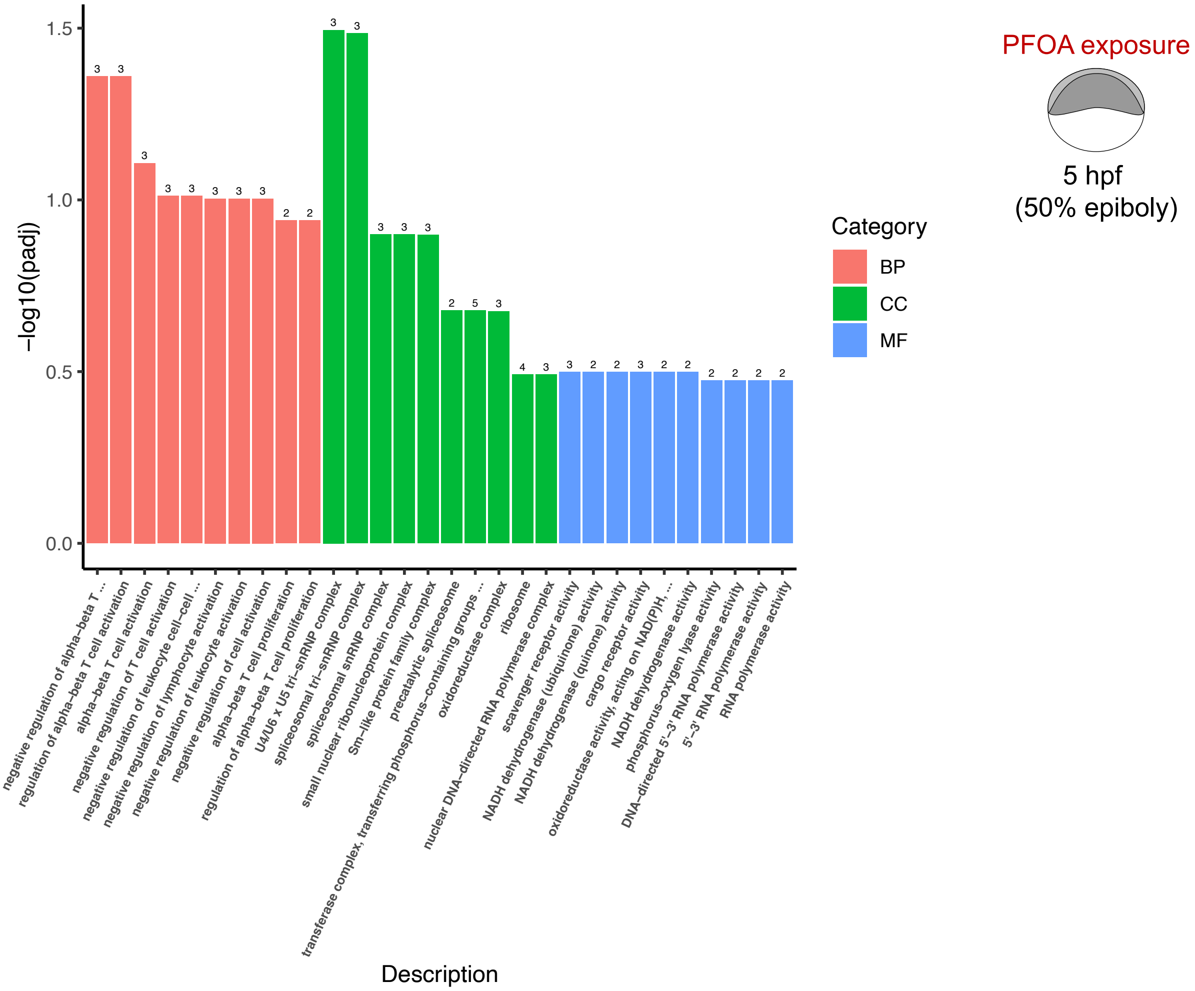


B) Kegg pathway analysis for PFOA exposure from 5 hpf compared against untreated

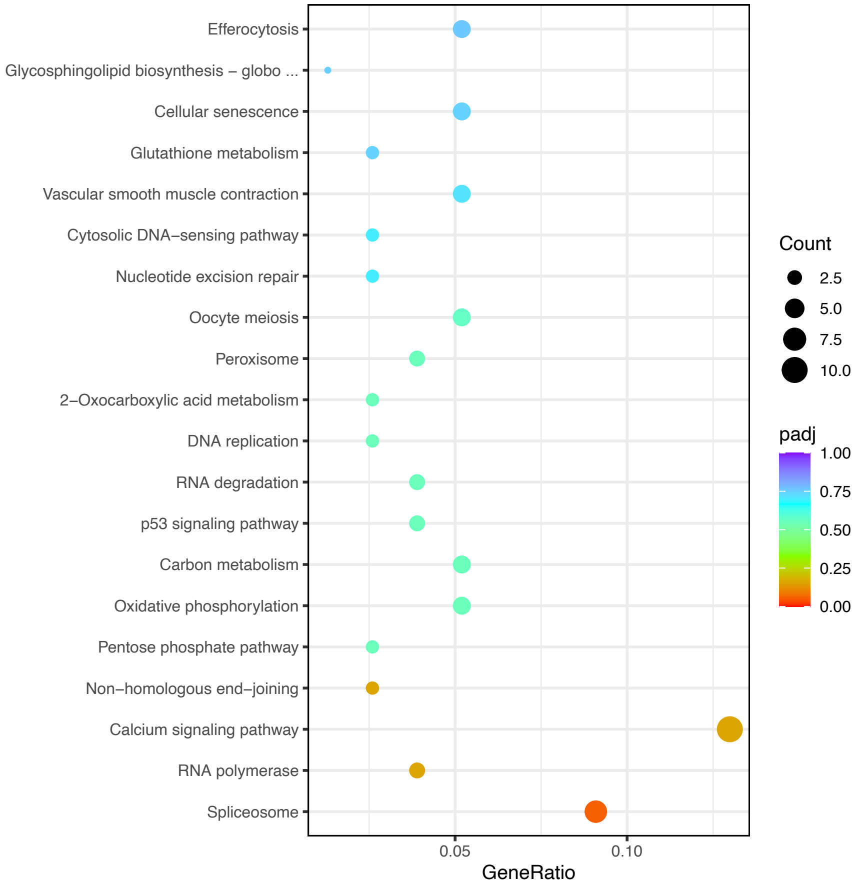

**Supp Figure 4**

A) GO term analysis for PFOA exposure from 8 hpf compared against untreated separated by categories

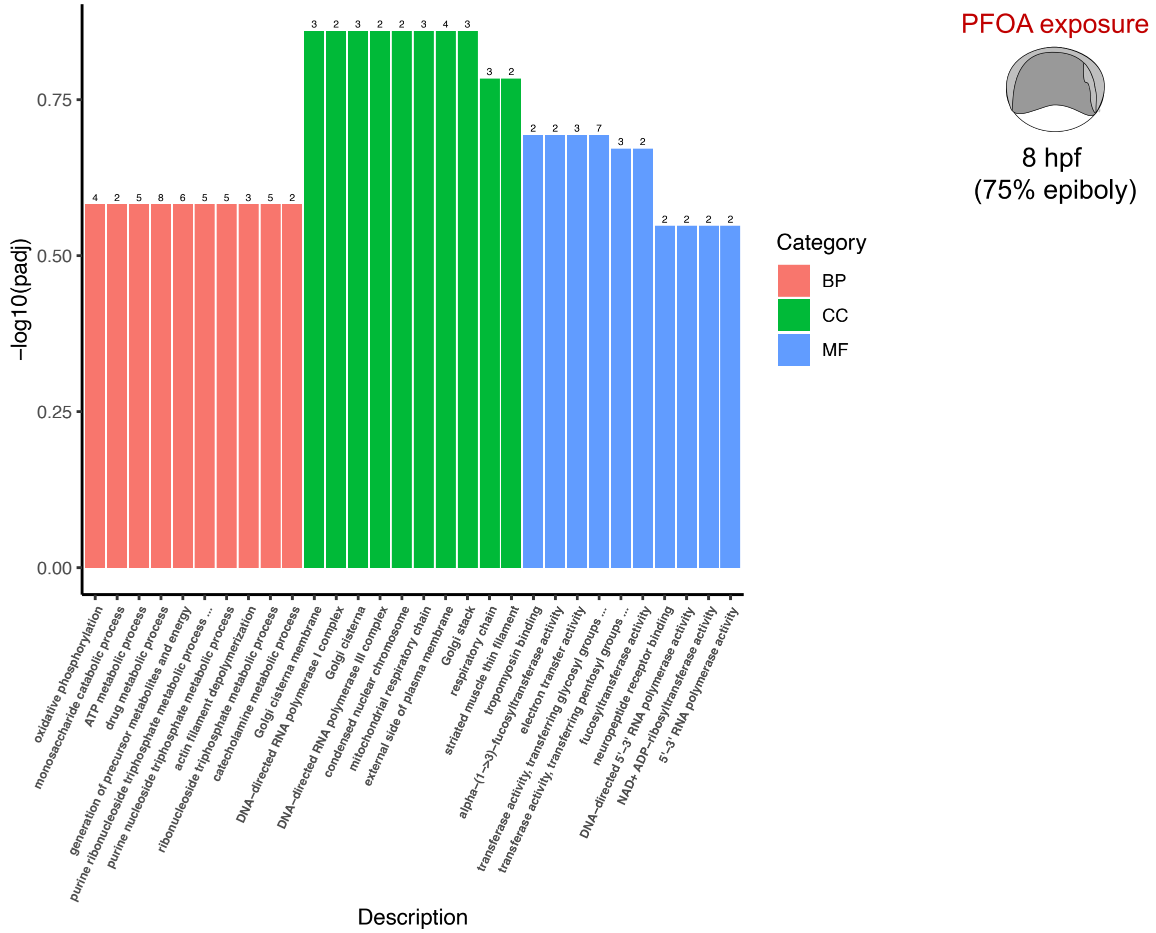


B) Kegg pathway analysis for PFOA exposure from 8 hpf compared against untreated

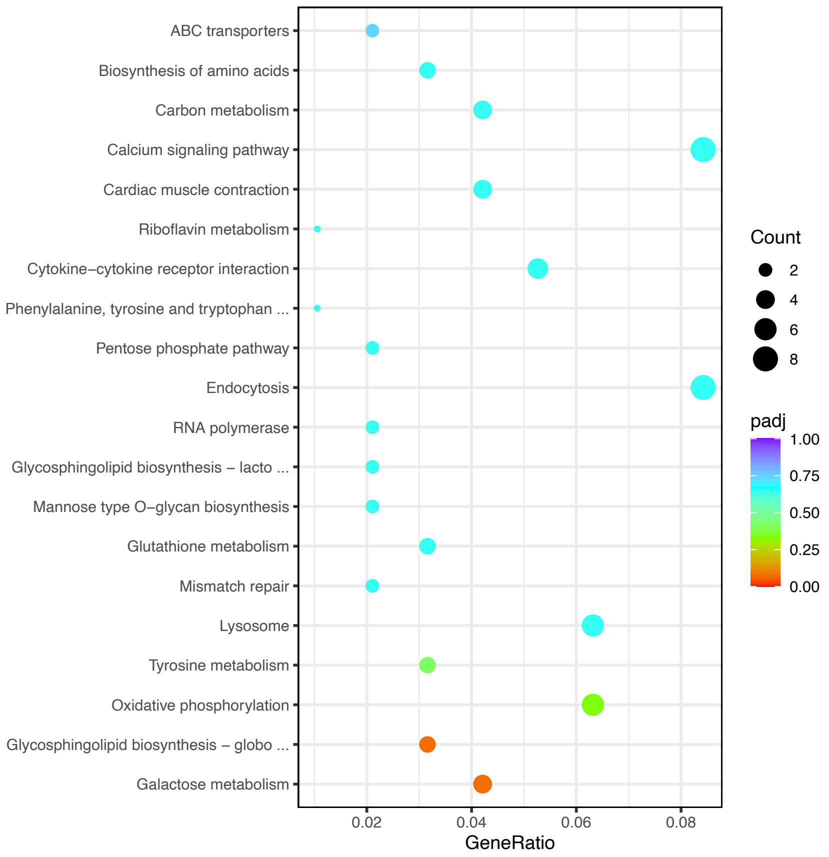

**Supp Figure 5**

Line graphs for Distance moved in Arena, Zone 1, and Zone 2 for different PFOA exposures

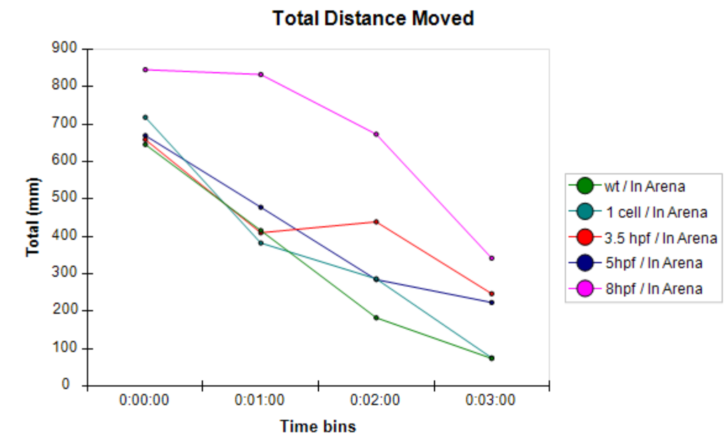


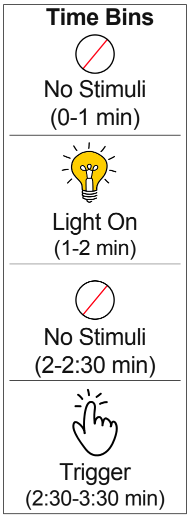


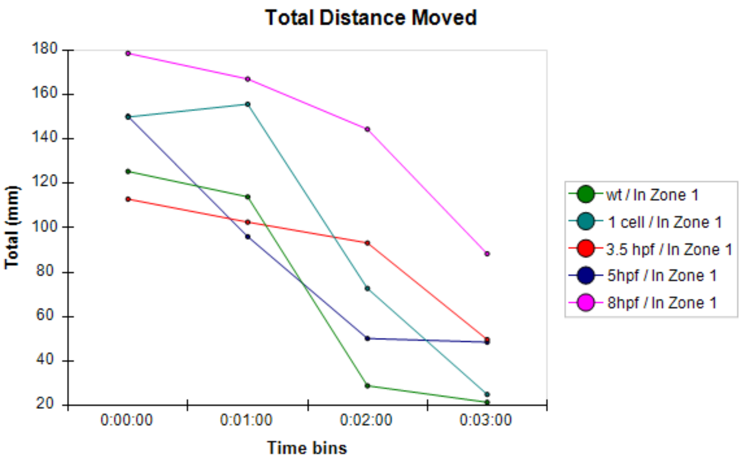


**Organization in 24-well plate**

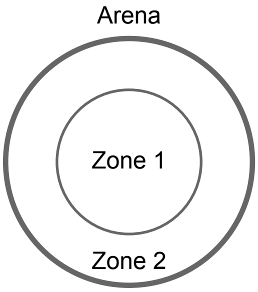


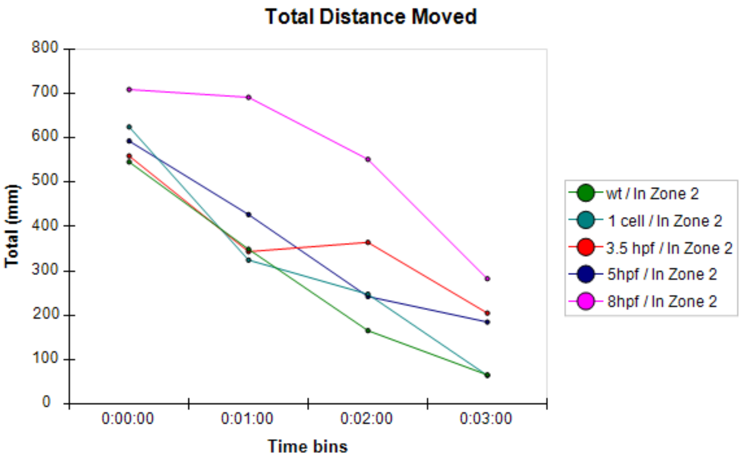


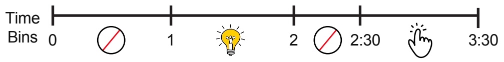


**Sample Name**

WT - Untreated

1 Cell - PFOA treatment from 1 cell

3.5 hpf - PFOA treatment from 3.5 hpf

5 hpf - PFOA treatment from 5 hpf

8 hpf - PFOA treatment from 8 hpf

**Supp Figure 6**

Line graphs for Velocity differences in Arena, Zone 1, and Zone 2 for different PFOA exposures

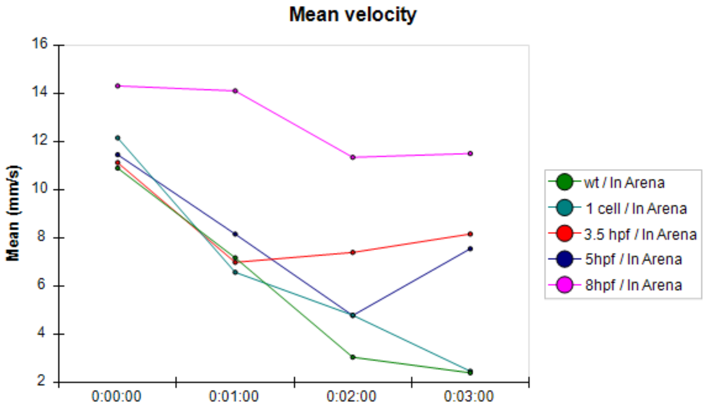


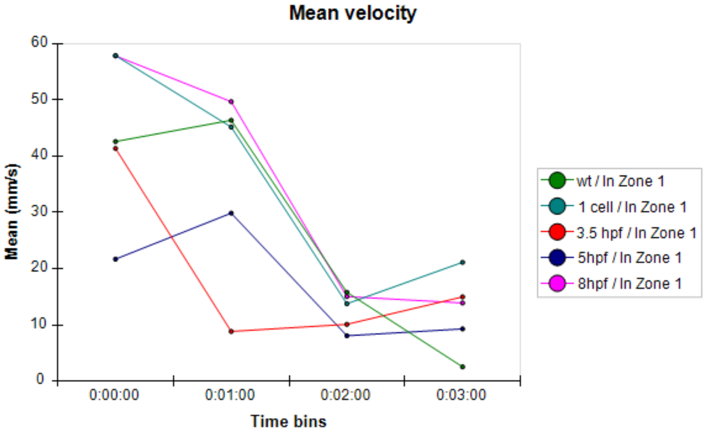


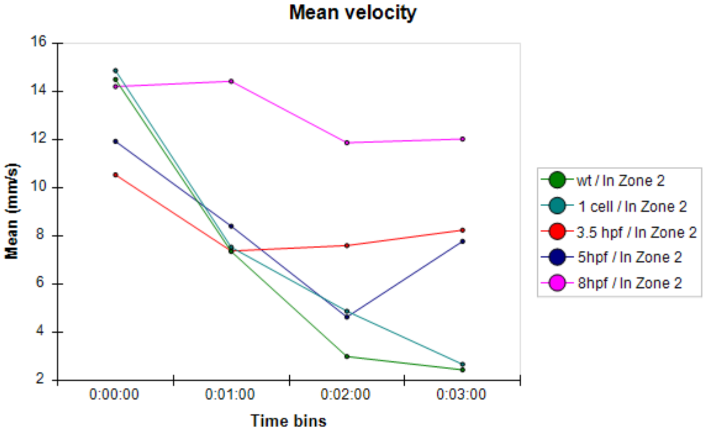


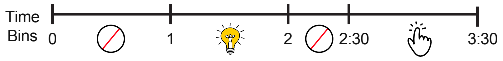


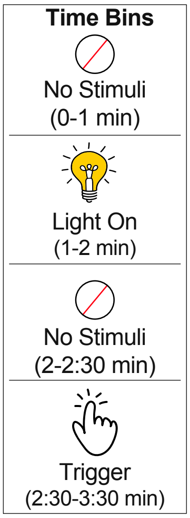


**Organization in 24-well plate**

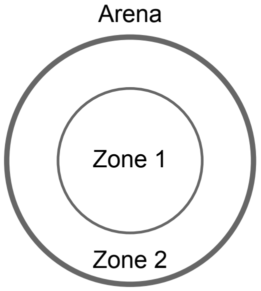


**Sample Name**

WT - Untreated

1 Cell - PFOA treatment from 1 cell

3.5 hpf - PFOA treatment from 3.5 hpf

5 hpf - PFOA treatment from 5 hpf

8 hpf - PFOA treatment from 8 hpf
